# Spanning-Tree Thermostatistics of Protein Allostery: An Exact Kirchhoff Framework with Application to Oncogenic KRAS

**DOI:** 10.64898/2026.04.29.721570

**Authors:** Fatma Senguler Ciftci, Burak Erman

## Abstract

This study introduces a statistical mechanical framework for allosteric communication in proteins based on the spanning-tree ensemble of residue contact networks. By representing *C_α_* protein backbones as weighted graphs, we identify each spanning tree as a topological microstate. The canonical partition function is evaluated analytically via the determinant of the reduced weighted Kirchhoff (Laplacian) matrix, allowing for the derivation of global thermodynamic functions (including Helmholtz free energy, internal energy, entropy, and heat capacity) without stochastic sampling.

Allosteric channels between specific residue pairs are defined as sub-ensembles containing unique simple paths. Using the Burton-Pemantle theorem and the Moore-Penrose pseudoinverse of the graph Laplacian, we compute path probabilities and channel-specific thermodynamics. This methodology enables a decomposition of channel heat capacity into energetic and topological components and quantifies residue-level allosteric importance through fractional contributions to the channel partition function.

The framework was applied to the G12D mutation in KRAS, comparing wild-type (PDB: 6GOD) and mutant (PDB: 6GOF) structures. Results show that while global thermodynamic properties remain highly conserved across the tight structural superposition, channel-level analysis shows a substantial internal redistribution of allosteric importance among intermediate residues, highlighted by the primary 12–61 signaling axis and distal routes (including shifts in residues such as Q61 and F156). Operating on *C_α_* backbone geometry, these topological shifts provide predictive hypotheses for subsequent molecular dynamics and experimental testing. Overall, this approach offers a rigorous, parameter-robust framework for understanding how point mutations perturb distal signaling networks.

## 1. Introduction

Allostery is the propagation of a regulatory signal between spatially distal sites in a protein. Two thermodynamic frameworks have traditionally interpreted this phenomenon. The first, due to Monod, Wyman, and Changeux [1], describes symmetry-breaking transitions between discrete conformational states. The second, due to Cooper and Dryden [2], attributes allostery to the entropic modulation of internal fluctuations without a change in mean structure [3-5]. Neither framework, however, accounts for the topological redundancy inherent in a protein’s contact network. A protein is not a collection of independent oscillators but a communication network whose signaling capacity derives from the multiplicity of available pathways.

Current structural models often employ local metrics, such as contact density, or path-based algorithms like betweenness centrality to identify allosteric hotspots [6-8]. More sophisticated approaches, such as eigenvector centrality based on mutual information [9], extend these ideas by considering the relative influence of a residue within the broader network. However, these methods remain essentially node-centric, focusing on the importance of individual residues rather than the collective thermodynamic properties of the signaling ensemble. Our framework shifts this perspective by treating the entire protein as a topological ensemble of spanning trees. This allows us to account for global coupling not as a ranking of influential nodes, but as a sum over all possible communication routes, enabling the calculation of channel-specific entropy and heat capacity.

This distinction is critical, as it bridges the gap between identifying influential nodes and defining the global landscape of the system. The central challenge of protein physics is thus to define a state function that quantifies this topological redundancy and identifies the specific bottlenecks where the inter-modular connectivity weakens, threatening the global integration of the network [10]. These integrated interactions suggest that allostery can be rigorously modeled as a topological problem using graph-theoretical methods. Indeed, graph-theoretical representations of proteins, where residues are treated as nodes and their physical interactions as edges, have become a cornerstone of structural bioinformatics [11-15]. Such models have been widely used to identify ‘small-world’ properties in protein folds and to locate high-centrality residues critical for stability. However, while standard graph methods often focus on static properties like shortest paths or degree centrality, they frequently overlook the full statistical complexity of the connectivity. By formalizing the protein as a contact network, we can employ exact combinatorial theorems to move beyond simple connectivity maps and toward a rigorous thermostatistical description of the allosteric state space. Central to this formalism is the conceptualization of the protein’s structural ensemble as a collection of cycle-free subgraphs known as spanning trees formed by the removal of just enough edges to eliminate all cycles while preserving global connectivity. Each such minimal subgraph contains exactly one route between any two vertices and possesses no redundancy. Different spanning trees represent distinct configurations of global connectivity, different ways a perturbation might propagate from one residue to another. The full protein graph, by contrast, contains many cycles. These cycles are the physical origin of alternative signaling routes. The set of all spanning trees of the graph constitutes a statistical ensemble. By identifying these spanning trees as the fundamental microstates of the protein graph, we can define the total signaling capacity through a canonical partition function [16].

By applying Kirchhoff’s Matrix-Tree Theorem [17], we obtain the partition function of the signaling landscape. This defines the topological free energy, which is the log-number of signaling routes, as its fundamental potential, and identifies the protein’s contact map not merely as a static structure, but as a generator for a canonical ensemble of acyclic microstates. A critical residue is one whose removal from the network substantially reduces the number of spanning trees, thereby contracting the ensemble of available signaling paths.

We applied the proposed formalism to the oncogenic G12D mutation in KRAS to investigate its constitutive activation. From the Burton–Pemantle probabilities [18, 19] and the Jensen–Shannon divergence [20] between the wild-type and mutant ensembles, the pathological effect of G12D emerges as a redistribution of residue centrality within the allosteric channels, while the channel thermodynamics, entropy, internal energy, and heat capacity, remain approximately constant. The global heat capacity remains robustly buffered (+0.50%), while the channels themselves maintain thermodynamic stability through strong internal compensation. This result identifies a precise thermodynamic signature of the G12D mutation: within individual channels, substantial but opposing shifts in energetic, topological, and cross-covariance components balance one another to conserve overall channel capacity. While this net channel capacity is conserved, the internal distribution of path occupancy across specific residues is heavily reorganized. This is evidenced by major shifts in allosteric importance at specific coordinate hubs (most notably a 35.5% redistribution at F156). Thus, the mutation does not alter the global fold capacity or the fundamental architecture of the channels, but instead systematically rewires how informational flux is routed through specific residue networks.

## 2. Theoretical framework

The framework in this section is developed in three stages. First, the thermostatistics of the global state is established by defining the spanning-tree ensemble for the entire protein graph. Second, the thermostatistics of individual channels is derived, mapping the global ensemble onto specific signaling routes between functional sites. Finally, an information-theoretic view of allostery is formulated, using the divergence between wild-type and mutant ensembles to quantify the change of signaling flexibility.

### 2.1 Global ensemble: thermostatistics of the protein graph

#### 2.1.1 Graph representation

The protein structure is represented as a weighted graph *G*(*V,E*), where the *N* vertices *V* denote *Cα* atoms and the edges *E* represent inter-residue contacts. The statistical mechanics of such a network is fundamentally governed by its connectivity.

#### 2.1.2 The topological microstate and cycle rank

The choice of the spanning tree as the fundamental microstate is rooted in the statistical mechanics of elastic networks [21], but its application extends to general information transfer. The signaling route is treated as a path along which a disturbance propagates [16]. In this view, the partition function sums all possible connected paths between residues, whether the coupling of the degrees of freedom between any two residues in the graph arises from vibrational modes, electrostatic shifts, or side-chain rotations. In Flory’s theory of phantom networks [21], the connectivity of an elastomer is characterized by its cycle rank *ξ* = *E* − *N* +1, which represents the inter-residue interactions beyond the minimal requirement for global connectivity.

Within this framework, the protein’s properties decompose into two physical origins. First, any individual spanning tree represents an acyclic microstate. Because its cycle rank is zero, it possesses no redundant constraints; its elasticity is purely intrinsic, defined by the sum of its edge weights. Second, the existence of cyclic edges, interactions in excess of the minimal spanning tree, generates topological entropy. These redundant interactions are the physical basis for correlations between alternative signaling paths. An interaction is defined as redundant if its removal does not disconnect the network but instead reduces its configurational diversity. This diversity is captured by the relationship between the number of spanning trees *τ* and the cycle rank *ξ* . While the removal of a redundant interaction decreases *ξ* by exactly 1, it can eliminate a disproportionately large number of spanning trees. Consequently, the capacity for information transfer is significantly decreased. The spanning-tree ensemble thus provides a discrete mapping that identifies information bottlenecks hidden within the global cycle rank, distinguishing between the mechanical presence of a bond and its actual contribution to the signaling ensemble.

#### 2.1.3 Edge weights and interaction parameters

To each edge *e*_*ij*_, a weight parameter *w*_*ij*_ is assigned. At an effective temperature kT (*in units of Å*), each edge receives a Boltzmann weight:

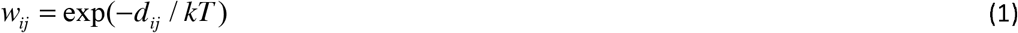

where *d*_*ij*_ is the inter-residue *C* _*α*_ −*C* _*α*_ distance. The weight is set to zero, *w*_*ij*_ = 0, whenever *d*_*ij*_ exceeds the contact cutoff *r*_*c*_, so that non-contacting pairs contribute no edge to the graph..

Physically, this exponential formulation is justified because the effective potentials governing non-bonded inter-residue interactions (such as van der Waals forces, hydrogen bonding, and hydrophobic packing) decay monotonically with distance. Treating the spatial separation *d*_*ij*_ as an intrinsic structural energy barrier ensures that tightly packed core residues contribute exponentially more to the allosteric signaling network than distal, poorly coordinated pairs.

Moreover, this formulation offers a significant physical improvement over classical Elastic Network Models (ENMs) and the Gaussian Network Model (GNM). While standard GNM relies on a binary step-function cutoff (treating all interactions within the cutoff identically and abruptly dropping to zero), our formulation introduces a continuous geometric decay. This eliminates unphysical step discontinuities at the cutoff boundary, prioritizing high-density structural conduits while preserving the exact determinantal tractability of the Kirchhoff framework. In this formulation, *d*_*ij*_ functions as the structural interaction parameter. The effective thermal scale *kT* is related to the physical temperature *k*_*B*_*T* through *kT* = *k*_*B*_*T* / *α*, where α is the contact energy parameter (units: energy/Å) that converts distance to contact energy. While α can be determined from experimental observables (such as crystallographic B-factors or NMR order parameters), no such calibration is pursued here. By adopting *kT* in units of Å, the physical force constant is absorbed into the temperature scale, rendering the Boltzmann exponent dimensionless. This mapping ensures that the statistical weights *w*_*ij*_ are determined solely by the protein’s geometry, and that the resulting thermodynamic state functions provide an unambiguous characterization of the differences between wild-type and mutant ensembles; throughout the paper, the heat capacity is expressed in dimensionless reduced units as the ratio of the physical heat capacity to the Boltzmann constant *C*^*^ = *C* / *k*_*B*_ .

No distinct weight is assigned to backbone (i, i+1) contacts; they are governed by Eq. (1) on the same footing as all non-bonded edges. It is nonetheless useful to ask what role the backbone plays within the resulting spanning-tree ensemble, since it is the sole structural element guaranteeing sequence connectivity. Using the marginal edge-inclusion identity *P* (*e* ∈*T*) = *w*_*e*_*R*_*e*_, where *R*_*e*_ is the effective resistance defined in Eq. (7) below, we find that backbone edges in KRAS (171 of the 796 edges of the wild-type contact graph, and 171 of the 803 edges of the G12D graph; the two graphs share 793 edges, the substitution gaining 10 contacts and losing 3 for a net increase of seven, see Section 3 and Table S4.1) are not topologically required for connectivity, none are bridges at the cutoff r_c = 7.8 Å used here, and in the unweighted (high-temperature) limit, are included in a random spanning tree at essentially their base rate (22% of expected tree edges, versus 21% of all edges). At the finite effective temperature used throughout this study (kT = 1.0 Å), however, backbone edges account for 58% of the expected edges in the weighted spanning-tree ensemble, a roughly ten-fold enrichment in mean inclusion probability relative to non-bonded contacts, consistently in both the wild-type and G12D structures. This behavior is an emergent consequence of Eq. (1): backbone bonds are uniformly short (∼3.8 Å) relative to the range of non-bonded distances admitted by the cutoff, so the exponential weighting privileges them without requiring a distinct parameter for the polymer backbone. This is consistent with the long-standing convention in the Gaussian Network Model of applying a single, uniform interaction weight to all contacts irrespective of bond type [15].

#### 2.1.4 Spanning trees as microstates

A spanning tree T of the protein graph *G* is a subgraph containing all *n* vertices connected by exactly *n* −1 edges. This configuration represents a state of ‘critical connectivity’. It is the minimal structure required to maintain global integration without the redundancy of cycles. The full protein graph, by contrast, contains a total of *E* edges; the difference *ξ* = *E* − *N* +1, i.e., the first Betti number [17, 21], quantifies the topological redundancy of the contact network. Within this framework, each tree serves as the combinatorial analogue of a normal mode: a collective, discrete state of the global interaction network. By identifying each spanning tree as a microstate, we effectively treat the allosteric signaling landscape as an ensemble of these non-redundant communication pathways.

#### 2.1.5 The global partition function

The canonical partition function Z is defined as the sum over all possible spanning trees:

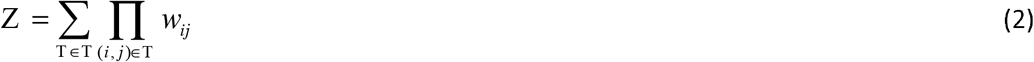

where T is a spanning tree and T is the set of all trees of the protein. By Kirchhoff’s Matrix-Tree Theorem [17], this sum is computed exactly by the determinant of the reduced weighted Laplacian matrix *L*_*w*_ :

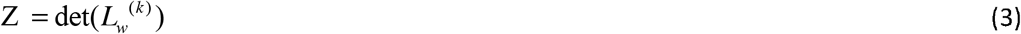

where 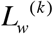 is the (n−1)×(n−1) reduced weighted Kirchhoff matrix obtained by deleting an arbitrary row t and its corresponding column t from the global Laplacian matrix *L*_*w*_ .

This identity defines the network free energy *F* = −*kT* ln Z . This framework recovers the classical Gaussian Network Model (GNM) [15, 22] in the high-temperature limit *kT* →∞, where all contact weights approach unity and Z becomes the number of spanning trees in the unweighted graph. A schematic representation of a protein graph, illustrating the relationship between spanning trees, redundant interactions, and topological entropy, is presented in Figure 1. As shown, even a simple 5-mer with a cycle rank of *ξ* = 3 generates an ensemble of 21 unique spanning trees. Although the following schematic assumes normalized edge weights (*w*_*ij*_ = 1) for clarity, in the general framework, each tree contributes to the global partition function Z according to its cumulative weight, defined as the product of its constituent edge weights *w*_*ij*_ .

**Figure 1:**
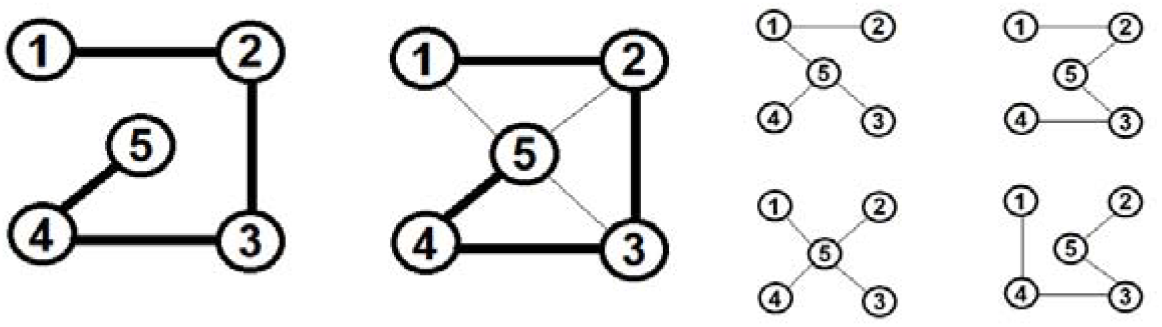
Graph-theoretic representation of a protein and its spanning-tree microstates. (Left) The primary sequence of a 5-mer is depicted as the simplest spanning tree, where heavy lines represent covalent peptide bonds. (Center) The folded state of the protein, where the topology is expanded by non-bonded contacts (thin lines). In this configuration, residue 5 acts as a hub connecting to residues 1, 2, and 3. This graph consists of 7 edges and possesses a cycle rank of *ξ* = 3, introducing redundant signaling pathways. (Right) A representative subset of four of the total 21 spanning trees that constitute the global communicative ensemble. Each tree represents a discrete, acyclic microstate of the network’s connectivity.

#### 2.1.6 The partition function and Kirchhoff’s theorem (revisited)

The energy *E*_*k*_ of a microstate *T*_*k*_ is defined as the sum of its constituent edge potentials:

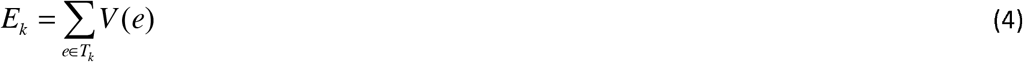

where *V* (*e*) = *d*_*e*_ is the contact potential assigned to edge e by Eq. (1), so that the microstate energy is simply the total contact length of the tree. According to the canonical formalism, the probability *P*(*T*_*k*_) of the system occupying a specific microstate is governed by the Boltzmann factor:

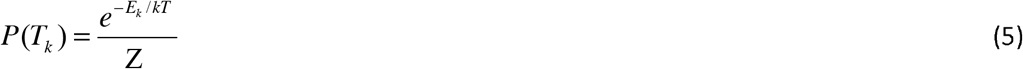

The Partition Function Z defined above by Eqs. 2 and 3 may also be represented as the sum over the manifold of all possible spanning trees:

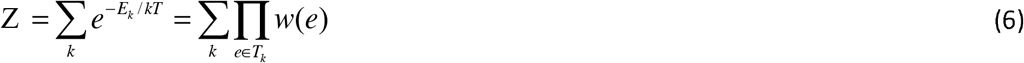

where *w*(*e*) = *e*^−*V* (*e*)/*kT*^ is the statistical weight of an interaction. Although the number of microstates scales exponentially with the number of residues, Kirchhoff’s Matrix-Tree Theorem permits the collapse of this sum into a single determinantal form given by Eq. 3. This result provides a direct mapping between the global topology of the protein and its thermodynamic state functions.

#### 2.1.7 Thermodynamic State Functions

From the partition function, we derive the fundamental thermodynamic quantities of the topological ensemble:

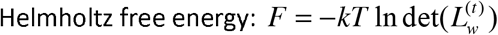

In this expression, *F* is measured in units of α, rendering *F* and *kT* dimensionally equivalent in Å.

Internal energy: 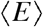

Entropy: 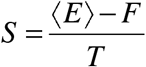

Heat capacity: 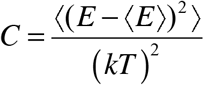

Here, averages are taken over all spanning trees and the Heat Capacity is in units of *k* and the averages are defined as 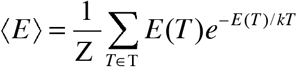 and 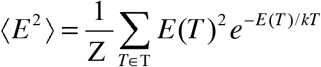.

The entropy *S* quantifies the configurational diversity of the communication network. A state of high entropy indicates a degenerate ensemble where numerous global signaling patterns coexist, whereas low entropy signifies a transition toward a rigid, deterministic structure. A peak in the heat capacity *C* indicates a cooperative reorganization of the dominant microstates, a combinatorial analogue of a phase transition.

### 2.2 Allosteric channel: local signaling between two residues

To understand allosteric communication, the protein must be viewed not merely as a single static configuration but as a distribution of states, where the signal is a collective property of the structural ensemble’s available microstates. These global properties are projected onto specific local channels to identify the most probable routes of information transfer using the Burton-Pemantle theorem [18, 19].

#### 2.2.1 Dynamic node-to-node distance

Before evaluating specific pathways, we must define the metric of connectivity within the ensemble. The foundation of our analysis begins with the Moore-Penrose inverse *K* ^†^ of the graph Laplacian *L*_*w*_ . We define the dynamic node-to-node distance *R*_*ij*_ between any two residues *i* and *j* as:

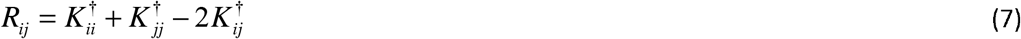

While the dynamic distance *R*_*ij*_ is calculated from a static crystallographic input (*d*_*ij*_), this framework is fundamentally grounded in physical fluctuation dynamics. In the context of Elastic Network Models, the static crystal structure is treated as the equilibrium minimum energy state (**R**^0^) of a multi-dimensional harmonic basin. Under the Gaussian Network Model (GNM) approximation, the potential energy *V* of the system due to thermal displacements Δ**R** is governed by the Kirchhoff matrix *L*_*w*_ as: 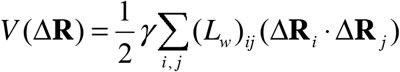 where *γ* is the single-parameter spring constant. Under the constraint of this harmonic potential, the system is immersed in a virtual thermal bath. Integrating over this multivariate Gaussian distribution yields the mean-square fluctuation of each residue: 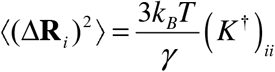. This equation of state provides the direct physical bridge to experimental crystallography, where the crystallographic B-factor measures the physical mean-square fluctuations of residues in the crystal lattice via 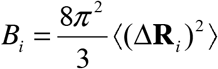. The well-documented, excellent correlation between the diagonal elements of the inverse Laplacian 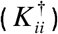 and experimental B-factors proves that the static network topology contains the exact physical constraints necessary to determine the amplitude and correlation of thermal fluctuations [15]. Thus, by taking the static crystal structure and calculating the generalized inverse of its Laplacian, we do not treat the protein as a rigid entity; rather, we solve the exact partition function of its harmonic fluctuations under the topological constraints of its folded state.

Physically, Eq. 7 represents the resistance to information flow between two points in the protein. Unlike a static Euclidean distance, the dynamic distance is a measure of the functional separation between two residues. A large dynamic distance indicates that the residues are topologically distant, meaning they possess high resistance to the coherent flow of information and lack the synchronized mechanical coupling required for signal propagation.

#### 2.2.2 Dynamic edge-to-edge distance

While residues are the coordinates of the protein, the signal propagates through the interactions (edges) between them. We therefore define the edge-to-edge dynamic distance matrix *Y*, which provides the mutual dynamic distance between any two edges *e*_*a*_ (connecting residues *i*_*a*_, *j*_*a*_) and *e*_*b*_ (connecting residues *i*_*b*_, *j*_*b*_):

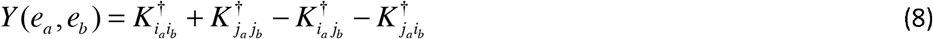

Physically, *Y* (*e*_*a*_, *e*_*b*_) quantifies the topological correlation between two interactions. It measures how much the mechanical load carried by one bond influences the requirement for another bond to be present within the same signaling ensemble.

#### 2.2.3 Path response matrix (Burton-Pemantle framework)

To quantify allosteric communication, we must transition from a node-based description of the protein to an edge-based ensemble of signaling routes. The probability that a given spanning tree contains a specific path *P* represents the frequency and reliability of that route across the entire topological ensemble.

For any specific signaling path *π* _*i*_ consisting of a set of *k* edges, we define a path-specific sub-matrix: the edge response matrix 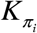. This *k* × *k* matrix is populated directly from the edge-to-edge dynamic distance values *Y*_*ab*_ defined in Section 2.2.2:

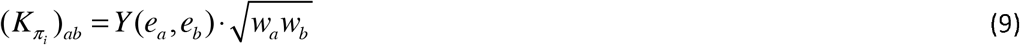

where *w*_*a*_ and *w*_*b*_ represent the weights (statistical weight) of the respective edges. The √(w_a w_b) prefactor serves as a normalization to render K_π dimensionless, rather than an independent reweighting of the edges. This matrix captures the internal correlations of the chosen path, reflecting how the individual interactions within that route are coupled to one another.

#### 2.2.4 Path probability via the Burton–Pemantle theorem

Following the Burton–Pemantle theorem [18, 19], the absolute probability *P*(*π* _*i*_) that all edges in the path *π* _*i*_ are simultaneously present in a weighted spanning tree is given by the determinant of the edge response matrix:

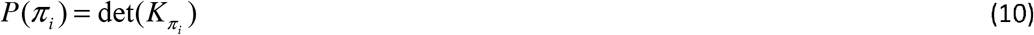

This determinant measures the functional volume of the configurational space occupied by the path *π* _*i*_ . If the edges in a path have a small edge-to-edge distance (indicating high synchronization), the determinant reflects the path’s high probability and dominance within the ensemble.

#### 2.2.5 Path occupancy and relative weights

Given a set of *m* simple paths {*π*_1_,*π* _2_,…,*π* _*m*_} identified between the endpoints, we normalize these absolute probabilities to compute the relative weights *w*(*π* _*j*_) :

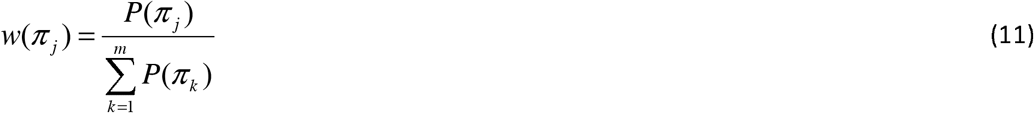

These weights represent the occupancy likelihood of each route. They demonstrate how the allosteric signal is distributed across the network’s available degrees of freedom.

#### 2.2.6 Topological coupling and the global entropic reservoir

A critical feature of the spanning tree ensemble is that an allosteric channel is never an isolated system. Although specific path sequences are enumerated, their relative statistical weights are governed by the global Laplacian matrix *L*_*w*_ . This creates a mathematical coupling where the remaining part of the protein acts as a topological heat bath.

To clarify how allostery is specifically associated with a group of spanning-tree microstates, we contrast our approach with traditional path-searching algorithms. In a single spanning tree (a topological microstate), there is exactly one unique, cycle-free path connecting any source residue *s* to any target residue *t* . A single tree therefore represents a single, non-redundant communication route. However, physical allostery is a collective network phenomenon operating over redundant, parallel pathways. We therefore define an allosteric channel as a sub-ensemble (or group) of spanning trees. Specifically, for a set of *M* possible simple paths 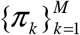 of a given length connecting *s* and *t*, each path *π* _*k*_ defines a distinct group of spanning trees, namely the set of all trees in the ensemble that contain the path *π* _*k*_ . The probability of this group of trees is determined exactly using the Burton-Pemantle theorem, mapping the local signaling route directly onto the global Moore-Penrose pseudoinverse *K* ^†^ . This definition allows us to treat the allosteric channel as a thermodynamic sub-ensemble. Under mutation, the total signaling capacity of this group of trees can remain thermodynamically stable while the system dynamically redistributes its microstate occupancies among the different alternative paths within the group. This models the protein’s ability to re-route signaling traffic in response to mutations without suffering a complete loss of distal communication.

The probability of any signaling path is not merely a product of its internal edge weights, but a measure of its compatibility with the global spanning tree ensemble. Every edge utilized by the channel restricts the possible configurations of the remaining residues to form a valid, cycle-free tree. This interaction creates an energy transfer between the channel and the surrounding protein.

This relationship is most visible in the path-associated heat capacity. Because the heat capacity represents the variance of energy fluctuations within the ensemble, the paths that maximize this expression are the protein’s most perturbable routes. These high-variance paths exist at the edge of topological stability, where the competition between diverse tree configurations is most intense. In an entropically open state (high configurational diversity), a high global heat capacity indicates a loose entropic reservoir. The background protein exerts low topological pressure, allowing the channel to explore a wide variety of path lengths and configurations. In a topologically constrained state (reduced configurational volume), a collapse in global heat capacity signals a transition toward a rigid topological background. This creates a direct topological coupling between the local signaling route and the global state of the protein manifold.

This entropic coupling explains how perturbations far from the primary signaling site can fundamentally change the allosteric response. By modifying the global Laplacian, a distant mutation or ligand binding event changes the topological background through which the signal must propagate. This background is defined as an entropic buffer: a collective reservoir of configurational freedom that allows the protein to absorb local fluctuations.

In an entropically open system, this buffer provides the topological slack necessary for the signaling channel to remain flexible and redundant. However, if the global reservoir is constrained, the buffer vanishes, and the resulting topological pressure forces the local signal into a rigid, deterministic state.

#### 2.2.7 Channel partition function and entropy

To quantify the signaling between *A* and *B*, we define the channel partition function, Z _*AB*_ . In a Boltzmann-weighted graph where each interaction *e* has a weight 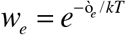, Z is the sum of the weights of all spanning trees that provide a unique path between these two residues. The channel free energy is then given by:

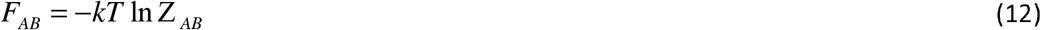

This potential represents the total topological resistance to signaling. A high *F*_*AB*_ indicates a channel that is either energetically costly or topologically sparse and signal propagation is difficult.

To characterize the internal structure of this channel, we define the path entropy *S* _*path*_ . This can be derived using two equivalent routes. First, as a Shannon entropy over the probability distribution of all possible signaling routes *π*_*i*_ :

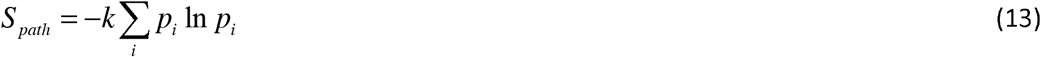

For any specific path *π*_*i*_ connecting *A* and *B*, its probability is defined as 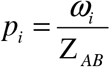, where *ω*_*i*_ is the sum of the weights of all spanning trees containing *π*_*i*_ . This represents the marginal probability of that path within the ensemble.

Alternatively, if we define the channel internal energy *U* _*AB*_ as the ensemble-averaged energy of all available paths *U*_*AB*_ = ∑ *p*_*i*_ *E*_*i*_, the entropy emerges from the fundamental thermodynamic relation:

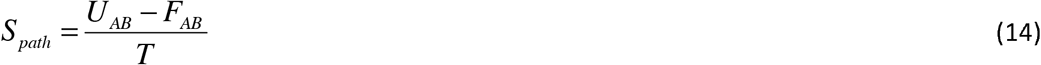

While the global internal energy of the protein is given by the ensemble average ⟨*E*⟩, we define the channel internal energy *U* _*AB*_ specifically for the sub-ensemble of paths connecting the allosteric and active sites.

The equivalence of these two formulations for the entropy confirms that allosteric communication is a balance between energetic favorability, low *U* _*AB*_, and configurational diversity, high *S* _*path*_ . If a single path dominates (*p*_*i*_ ≈ 1), *S* _*path*_ vanishes, indicating a rigid, non-redundant signaling bottleneck. If many degenerate paths coexist, *S* _*path*_ is maximized, providing the protein with high mutational robustness and the ability to tune its response through subtle shifts in the ensemble.

#### 2.2.8 Channel Heat Capacity and Route Flexibility

The final signature of the allosteric ensemble is the channel heat capacity, *C*_*AB*_ . Following the classical canonical ensemble formulation, the channel heat capacity is defined as the derivative of the channel internal energy (*U* _*AB*_) with respect to temperature:

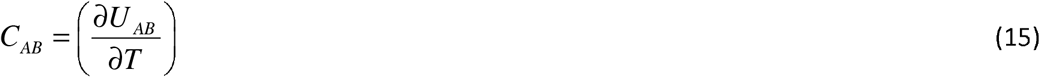

In our topological framework, *C*_*AB*_ is directly proportional to the variance of the effective energy states sampled along the path ensemble within the channel. Mathematically, it is expressed as:

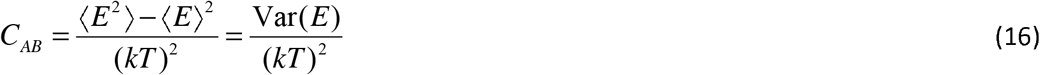

where *C*_*AB*_ is given in dimensionless units of the Boltzmann constant *k* . This variance represents the total spread or diversity of the path ensemble. A high channel heat capacity indicates that information is distributed across many parallel pathways with distinct weights, whereas a low heat capacity suggests that the ensemble is concentrated into a few dominant structural routes.

To show that the constituent contributions to this capacity are mathematically unique, rigorous, and completely non-arbitrary, we derive a formal decomposition directly from the fundamental theorem of probability for the variance of a sum of random variables. For any spanning tree microstate within the channel ensemble, its total effective energy (*E*) is defined as the sum of two distinct physical contributions:

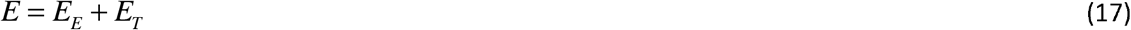

where *E*_*E*_ is the physical path distance (the sum of the physical edge lengths along the path) and *E*_*T*_ is the topological contribution of the surrounding network.

Applying the algebraic variance identity for the sum of two real-valued random variables, Var(*A* + *B*) = Var(*A*) + Var(*B*) + 2Cov(*A, B*), to Equation 17 yields:

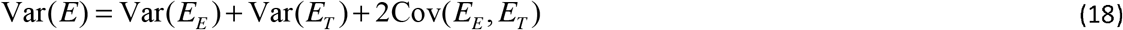

Dividing this exact identity by (*kT*)^2^ produces the precise, unique decomposition of the total channel heat capacity presented in our study: *C*_*AB*_ = *C*_*AB,E*_ + *C*_*AB,T*_ + *C*_*AB, X*_ where the three constituent thermodynamic terms are defined as follows:

Physical Path Component, 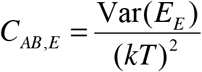. This term quantifies the thermal fluctuations in the cumulative physical length of the channel, reflecting the geometric flexibility of the local bonds that define the path.

Topological Path Component, 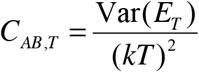. This term measures the fluctuations in topological slack, accounting for the variance in spanning-tree degeneracy. It represents the energy transferred from the surrounding protein’s configurational reservoir as it maintains global structural connectivity.

Cross-Coupling Component, 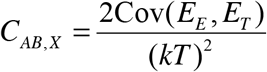. This term captures the dynamic correlation between the local physical fluctuations and global topological reorganizations.

The covariance term is physically vital to the allosteric mechanism. When a local path length (*E*_*E*_) fluctuates due to thermal motion, it shifts the overall network constraints, which in turn alters the entropic reservoir of available spanning tree configurations (*E*_*T*_) in the surrounding matrix. Without this cross-term, the mathematical representation of this local-to-global feedback loop would be entirely lost, and the individual components would fail to sum to the total channel heat capacity.

### 2.3 Information theory and mutational perturbation

#### 2.3.1 Information-theoretic response to mutation

A mutation at residue *r* is treated as a perturbation of the weighted Laplacian, *L*_*w,mut*_ = *L*_*w,wt*_ + Δ*L*_*w*_ . Because all thermodynamic quantities are derived directly from the Laplacian matrix *L*_*w*_, the impact of a mutation is calculated exactly, rather than estimated. The effect of a single change at a distant site propagates through the determinant of the matrix, which naturally accounts for the entire protein structure at once. This allows us to observe the global shift in the signaling ensemble without relying on simplified guesses or manual approximations.

This shift is quantified by the Kullback–Leibler Divergence *D*_*KL*_ :

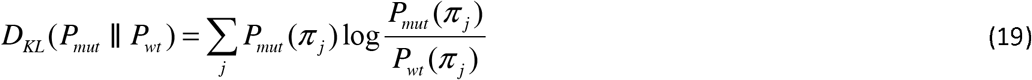

A significant *D*_*KL*_ signifies a fundamental reorganization of the allosteric ensemble.

To ensure a symmetric and bounded comparison between the wild-type *P* and mutant *Q* distributions, we further utilize the Jensen-Shannon Divergence *D*_*JS*_ [20]:

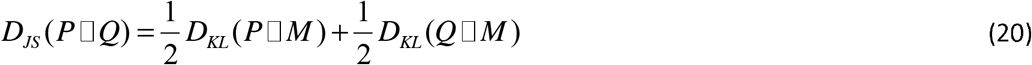

where 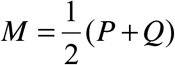 is the mixture distribution. While *D*_KL_ identifies the directional shift in information, the *D*_*JS*_ provides a normalized measure of the overall similarity between ensembles. A negligible *D*_*JS*_, as observed in our results, indicates that the global informational capacity of the channel is preserved, even when specific internal paths are significantly re-partitioned.

#### 2.3.2 Participation ratio and allosteric importance

To characterize how mutations reshape these networks, we define the following parameters, all of which are derived from the global cycle rank *ξ* = *E* − *N* +1:

1. Participation ratio *PR*_*AB*_ : Defined as 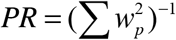, this is the exponential of the second-order Rényi entropy *H* _2_ [23]. It characterizes signal localization, providing a measure of the effective number of paths contributing to the allosteric channel. PR ranges between 1, corresponding to the signal being concentrated on a single path, and N, corresponding to the signal being distributed uniformly across all N paths. A low PR signals a transition toward a singular, dominant communication route, a hallmark of mutational bottlenecking.
2. Allosteric importance *I* of a residue k: 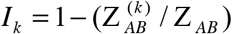, identifying bottleneck residues whose mutation most significantly disrupts the channel partition function. Here, 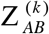 is the constrained partition function that specifically counts the signaling pathways between two distant points (Residues *A* and *B*) that must pass through residue *k* .

To prevent any conceptual overlap with classical graph-theory centralities, we explicitly distinguish Allosteric Importance (*I*_*k*_) from standard betweenness centrality and global structural importance.

First, unlike shortest-path betweenness centrality, which is a static, temperature-independent metric restricted to geodesic paths, *I*_*k*_ is a thermodynamic state function. It is evaluated over a canonical ensemble of spanning trees and accounts for all possible self-avoiding paths connecting two sites, weighting them physically according to their Boltzmann factors. This introduces physical temperature dependence, allowing the signal to shift from highly localized paths at low temperatures to diffuse, high-entropy pathways at physiological temperatures.

Second, *I*_*k*_ distinguishes targeted allosteric importance from global structural importance. While a node’s structural importance is a global measure of its role in overall network cohesion, *I*_*k*_ is intrinsically boundary-dependent. It is defined strictly relative to a specified Source (effector site *A*) and Target (active site *B*). A residue with low global connectivity can therefore exhibit high *I*_*k*_ if it serves as a critical bottleneck for that specific channel.

Finally, this formulation does not require any prior knowledge of the allosteric pathways. The only required inputs are the two physical endpoints *A* and *B* . The Kirchhoff framework automatically solves the partition function over all intermediate routes using the Burton-Pemantle theorem, meaning that the allosteric pathways are not assumed beforehand, but are instead discovered mathematically through the emergence of high-*I*_*k*_ residue peaks.

These functions, participation ratio PR and allosteric importance *I*_*k*_, are the statistical consequences of the redundant interactions within the protein graph. In the limit where the cycle rank is zero, which physically corresponds to stripping away all three-dimensional, non-bonded tertiary contacts to leave only the covalent polypeptide backbone (the linear primary chain), the network collapses into a unique spanning tree. Because this singular backbone-only microstate offers no alternative signaling paths or configurational diversity, the partition function *Z* reduces to unity, the topological entropy drops to zero (*S* = *k*_*B*_ ln(1) = 0), and all topological response functions (such as the topological heat capacity) vanish.

#### 2.3.3 The convergence test: dynamic distance as a sum-rule

A fundamental challenge in the characterization of the channel is the combinatorial explosion of possible paths as the path length increases. For a complex protein graph, the number of self-avoiding paths grows exponentially, rendering an exhaustive summation of the partition function computationally intractable beyond a threshold, typically lengths of 7 to 9 residue paths. To resolve this, we treat the channel as a truncated ensemble.

To evaluate the signaling channel between nodes *a* and *b*, we first calculate the exact effective resistance *R*_*ab*_ from the full Kirchhoff inverse of the global Laplacian. This value represents the ground truth of the entire protein’s connectivity. We then select a finite subset of paths from a to b as our local ensemble and construct the sparse matrix, *L*_*w,subset*_, containing only the nodes and edges that exist within our chosen ensemble of signaling paths. Empirical validation on different systems shows that channels constructed by paths of node length 9 captures more than 93-99% of the total information flow defined by the full matrix. This convergence proves that allosteric communication is dominated by a high-probability core of relatively short paths, and that the truncated Shannon entropy provides a numerically exact lower bound for the true functional diversity of the channel.

#### 2.3.4 The principle of minimum surprisal

The allosteric channel is governed by a partition function *Z*_*AB*_ . Since every spanning tree contains exactly one unique simple path between two nodes, the set of all possible paths is exhaustive and mutually exclusive over the tree ensemble. Therefore, the sum of their raw probabilities must equal 1

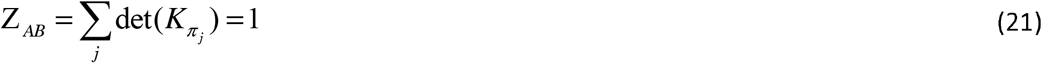

The Boltzmann path distribution thus simplifies to

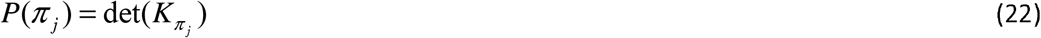

To quantify how favored a specific route is within the protein, we calculate its surprisal *I* (*π* _*i*_) . Mathematically, this is the negative logarithm of the path probability,

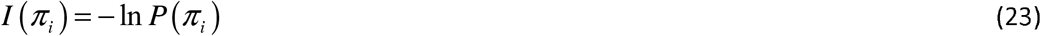

Physically, we can interpret this as the signaling potential: a low surprisal path means that the channel is a natural highway. It has a high probability because it is both energetically cheap and topologically accessible. The protein prefers this route. A high surprisal channel, on the other hand, means that the path is a rare detour. It has a low probability because it is either blocked by weak contacts or masked by more efficient alternatives.

Substituting from 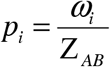

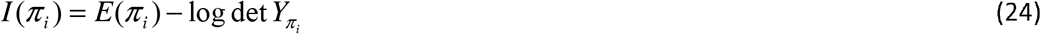

Here, *E*(*π* _*i*_) is the bare energetic cost of the path, representing the total topological resistance encountered by a signal traversing that specific route. A higher cost indicates a path comprised of weaker or less frequent inter-residue contacts, making it a less efficient channel for allosteric communication. log det 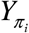 represents a topological bonus. This bonus quantifies the degree to which the surrounding network redundancy reduces the surprisal of a path below its intrinsic energetic cost. The global diversity of the communication routes is measured by the channel entropy *S*_*AB*_ given by Eq. 13.

Following the definition of the channel heat capacity, we propose that the functional state of a protein is one that minimizes the self-information *I* (*π* _*i*_) of the signaling ensemble. Just as a system in thermal equilibrium minimizes its Helmholtz free energy, an allosteric signal selects the routes that maximize the topological bonus log det 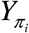.

This allows us to identify the dominant allosteric mode as the path (or set of paths) that satisfies the extremum of the probability distribution 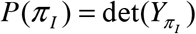. Any structural perturbation that increases the surprisal of the channel effectively weakens the communication, acting as an increased resistance to the flow of biological information.

Rather than assuming an evolutionary or functional optimization process, the path distribution in our model is governed by a standard statistical mechanical variational principle. In the canonical ensemble of spanning trees, the probability *P*(*π*_*i*_) of any given signaling path *π*_*i*_ is determined by the joint contributions of its local physical interactions and its global topological integration. Under the Burton-Pemantle theorem, this path probability is mathematically defined as:

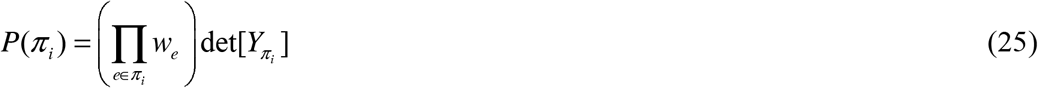

where the prefactor 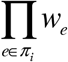 represents the bare energetic weights (statistical weight) of the constituent edges along the path, and 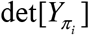 is the determinant of the edge-to-edge dynamic distance matrix, reflecting the topological correlations imposed by the surrounding cyclic network.

By taking the negative logarithm of this probability, we obtain the path’s self-information (or surprisal), *I* (*π* _*i*_) = −ln *P*(*π*_*i*_) . This formulation decomposes the surprisal into two physically distinct terms:

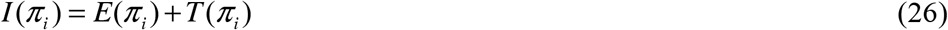

where 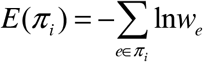 represents the cumulative energetic resistance of the path, and 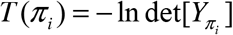 represents the topological correlation cost.

Under thermal equilibrium, the signaling ensemble naturally distributes its communication density to maximize probability, which is mathematically equivalent to minimizing the path surprisal *I* (*π*_*i*_) . Consequently, the dominant allosteric channels are identified as those that satisfy this joint minimization. This extremum does not depend solely on the topological determinant; rather, it balances the bare energetic cost of the contacts against the cooperative topological bonus offered by the surrounding network structure.

To address the combinatorial explosion of self-avoiding paths in highly cyclic protein networks and ensure the mathematical convergence of the channel partition function, we implement a localized, thermodynamics-driven path truncation protocol. The selection of active signaling paths is governed by the following dual physical criteria:

First, we compute the global effective resistance *R*_*ab*_ analytically using the Moore-Penrose pseudoinverse of the Kirchhoff Laplacian matrix (*L*^+^). Because this calculation is exact and does not rely on path enumeration, it provides an absolute, unapproximated physical baseline for the total topological connectivity between residues *a* and *b* .

Second, we restrict the graph traversal to a maximum path length of 9 residues. This spatial constraint defines our localized allosteric envelope, filtering out diffuse whole-molecule network noise and preventing combinatorial overflow. Within this search envelope, we exploit the exponential decay of path probabilities with increasing physical length. The probability *P*(*π*_*i*_) of any given path *π*_*i*_ is determined by its cumulative structural energy:

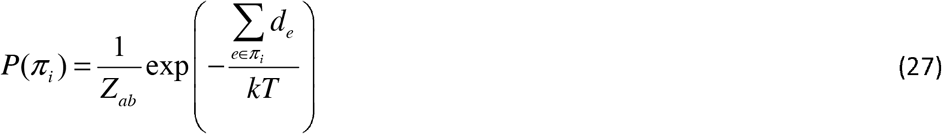

Due to this Boltzmann-like scaling, long paths or those traversing weak contacts have extremely high cumulative distances, resulting in exponentially vanishing probabilities that represent negligible thermal noise. We systematically sort all paths within the length-9 envelope in ascending order of their surprisal, *I* (*π* _*i*_) = −ln *P*(*π*_*i*_) . We then construct the active channel sub-ensemble P_active_ by retaining the top-ranking paths that satisfy a cumulative probability threshold *η* :

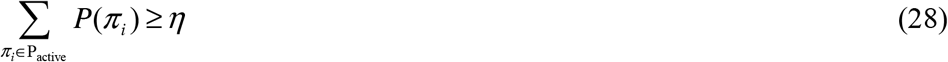

where we set *η* = 0.99 to capture 99% of the communication probability space contained within the corridor.

Finally, the convergence of this active subset is validated by verifying that the effective resistance reconstructed solely from P_active_ matches the exact analytical ground truth *R*_*ab*_ within a strict relative tolerance of ò < 10^−3^ . Sensitivity audits extending this traversal to a path length of 10 yield negligible variance in the calculated thermodynamic properties, (See Supplementary Material, Section S1, Table S1.1) confirming that a 9-residue limit represents the exact physical boundary where localized allosteric information is maximized and computational tractability is strictly maintained.

### 2.3.5 Distribution of the information current and signal convergence in the allosteric channel

The negative correlation proven by the Burton-Pemantle theorem (Eqs. 10 and 11) refers to the occupancy of edges. If one interaction (edge) is included in a signaling tree, it reduces the statistical likelihood that a competing interaction will be included in the same spanning tree. This mathematical property dictates a strict interdependence among the members of the spanning-tree ensemble: since the sum of all path probabilities is exhaustive and equal to 1, any change in the weight of one path must be balanced by the others.

In the language of information theory, we liken this channel to a discrete memoryless system where the alphabet consists of the available topological routes. If a specific trajectory is topologically disrupted (its weight *w*_*ij*_ decreases), the probability mass, which is defined as the information current, is necessarily redistributed across the remaining members of the alphabet. Conversely, if a mutation energetically strengthens a single route, the negative correlations require that it funnels the information away from the rest of the ensemble.

While the previous paragraph established the mathematical interdependence of the entire network, allosteric regulation specifically requires the transmission of a signal between distant functional sites. The focus now shifts from the global ensemble to the specific channel that connects these coordinates, applying information theory to quantify how the protein’s topology dictates signal reliability.

#### 2.3.6 Definition of the communication channel

While the spanning-tree ensemble {*T*_*k*_ } characterizes the global state of the protein, allosteric regulation is defined by the flow of information between a source residue *s* and a target residue *t* . We define the allosteric channel C(*s,t*) as the sub-ensemble of all simple paths {*π* _*j*_ } connecting these two coordinates.

## 3. Mutation effects on KRAS

We applied this framework to KRAS, a key player in cell growth and one of the most common drivers of human cancers [24]. We sought to answer a specific question: Does the G12D mutation create entirely new signaling pathways, or does it use the same residue paths but change how often each path is actually used?

We analyzed two KRAS crystal structures in their active, GNP-bound conformations: the wild-type (PDB code: 6GOD) and the G12D mutant (PDB code: 6GOF). Both structures yield 172 residues in chain A. For each protein, *C* _*α*_ contact graphs were constructed using a cutoff of *r*_*c*_ = 7.8 *Å* to define the interaction topology. From these graphs, global thermodynamic properties (energy, entropy, and heat capacity) were evaluated analytically using the graph Laplacian spectrum via Kirchhoff’s Matrix-Tree Theorem rather than via stochastic sampling [17, 25].

The overall network topologies of 6GOD and 6GOF remain exceptionally similar due to their tight structural superposition (Least-squares superposition of all 172 Cα atoms) of chain A of 6GOF onto the corresponding atoms of 6GOD (one-to-one residue correspondence, no trimming) gives an RMSD of 0.19 Å and a TM-score of 0.998; the largest single deviation, 0.98 Å, occurs at the N-terminal residue. The G12D substitution therefore produces no detectable backbone rearrangement in the crystal, and the two contact graphs are correspondingly similar: of the 806 residue pairs in contact in at least one structure, 793 are shared (Jaccard overlap 0.984), giving 796 contacts in the wild type and 803 in the mutant at *r*_*c*_ = 7.8 Å. Only 13 pairs differ, 10 present exclusively in G12D, 3 exclusively in the wild type, a net gain of seven edges, and all 13 lie between 7.5 and 8.2 Å in both structures. These are contacts sitting marginally on either side of the cutoff rather than the signature of a conformational change, and no contact involving residue 12 itself crosses the threshold. Although pairs such as F28–T148 and G77–E162 are remote in sequence, the tertiary fold places them in close spatial proximity (*d*_*ij*_ ≈ 7.7 Å in 6GOF), so sub-Ångström backbone adjustments are sufficient to open or close these local bridges and thereby to alter path counts in the distal channels (Supplementary Section S4, Table S4.1 and Figure S4.1). Minor path count variations in distal channels simply reflect subtle inter-residue contacts crossing the cutoff rather than global conformational changes (see Supplementary Material S4, Table S4.1, and Figure S4.1).

A key strength of this framework is that the global partition function is evaluated analytically without stochastic sampling error. Nevertheless, the implementation relies on defined input parameters, including the spatial contact cutoff (*r*_*c*_ = 7.8 *Å*), effective thermal energy (kT), and a truncated channel expansion. To confirm the robustness of our results, we evaluated parameter sensitivity across cutoffs of 7.5-8.5*Å* and thermal energies of *kT* = 0.8-1.2 . The relative channel shifts and primary residue contributions remain highly stable across this parameter space (Supplementary Note S5, Figure S5.1), demonstrating that the observed topological response is a robust physical signal.

In summary, because our model relies on *C*_*α*_ backbone coordinates, it detects the effect of the mutation through backbone geometric adjustments rather than explicit side-chain chemistry. Thus, the residue-level path shifts identified here serve as predictive hypotheses, with all-atom Molecular Dynamics (MD) or experimental testing (e.g., mutagenesis or NMR) representing natural next steps for validation.

### 3.1 Global Thermodynamics

The global thermodynamic properties of the wild-type and G12D systems, derived from the spanning-tree partition function at a reduced temperature *T* = 1.0, are summarized in Table 1. At the global scale, the mean energy ⟨*E*⟩ (+0.13%) and entropy *S* (+0.37%) remain remarkably consistent between the two states. Furthermore, the global heat capacity *C* exhibits a negligible increase from 139.6 in the wild type to 140.3 in the G12D mutant, a shift of just +0.50%.

**Table 1.** Global Thermodynamics of WT and G12D KRAS.

| Property | WT | G12D | % |
| --- | --- | --- | --- |
| Energy, $\langle E \rangle$ | 783 | 784 | +0.13 |
| Entropy, $S$ | 267 | 268 | +0.37 |
| Heat capacity, $C$ | 139.6 | 140.3 | +0.50 |
| Number of nodes, $N$ | 172 | 172 | 0.00 |

*Note.* $\% = 100 \times (G12D - WT)/WT$ .

This remarkable invariance demonstrates that KRAS is a structurally stable fold whose overall thermodynamic landscape is robustly buffered against single-site oncogenic mutations. The mutation does not induce global rigidification, structural collapse, or dramatic landscape narrowing. However, this global stability masks critical localized dynamics: mean thermodynamic metrics across the entire spanning-tree ensemble cannot show how informational flux is redistributed internally.

To determine how the oncogenic mutation alters signal transmission without disrupting global fold thermodynamics, we must transition from a global ensemble average to a channel-resolved examination of individual allosteric routes.

### 3.2 Channel thermodynamics

Utilizing the Burton–Pemantle theorem as detailed in Sections 2.2.3 and 2.2.4, we conducted an exhaustive enumeration of path probabilities within the wild-type (WT) and G12D KRAS ensembles. Rather than evaluating isolated trajectories, our analysis identifies ten primary signaling channels (*L* ∈[2, 9]) that define the major thermodynamic corridors of information transfer across the protein structure.

These ten channels capture the multi-scale allosteric architecture of KRAS and are categorized into four functional groups:

1. **Primary Switch-Direct Axis (Channels 6, 7, 10)**
  - **Channel 7 (Residues 12** → **61; G12**→**Q61/SwII):** The canonical signaling axis connecting the P-loop directly to the catalytic Switch II region. It represents the primary thermodynamic anchor where informational flux encounters minimal topological resistance.
  - **Channel 6 (Residues 12** → **35; P-loop**→**SwI):** Connects the P-loop directly to the Switch I loop, mediating nucleotide-dependent effector engagement.
  - **Channel 10 (Residues 35** → **61; SwI**→**SwII):** Represents direct inter-switch coupling, governing the coordinated conformational response between Switch I and Switch II.
2. **Active-Site Flanking Buffers (Channels 1 and 2)**
  - **Channel 1 (Residues 6** → **11; P-loop):** Located entirely within the P-loop (*β*1-strand), flanking the active site. It monitors local phosphate-binding dynamics.
  - **Channel 2 (Residues 55** → **60; pre-Switch II):** Positioned on the *β* 3 -strand, bridging nucleotide-sensing and effector-recognition loops as a flexible hinge.
3. **Nucleotide Recognition and Inter-Lobe Corridors (Channels 3, 4, and 5)**
  - **Channel 3 (Residues 110** → **117; GBS):** Located near the Guanine Binding Site (GBS), sampling allosteric responses driven by base recognition.
  - **Channel 4 (Residues 141** → **146; SAK motif):** Part of the SAK motif on *β* 6, monitoring long-
  - range topological pressure propagated to the distal nucleotide-binding interface.
  - **Channel 5 (Residues 19** → **142; Lobe Linker):** The primary inter-lobe bridge connecting Lobe 1 (residues 1–86) and Lobe 2 (residues 87–166) via the L19–I142 hydrophobic contact.
4. **Distal C-Terminal Extensions (Channels 8 and 9)**
  - **Channel 8 (Residues 12** → **156; P-loop**→*α* 5 **/** *β* 6 ): Connects active-site fluctuations to the C-terminal *α* 5 -helix/ *β* 6 -sheet structural anchor.
  - **Channel 9 (Residues 12** → **170; P-loop**→**C-term):** Extends across the longest structural axis to the C-terminal hypervariable boundary.

Together, these ten channels capture the entire allosteric manifold: the switch-direct channels provide primary signal transfer, while the flanking, inter-lobe, and distal corridors supply the topological slack necessary to buffer or redirect thermal fluctuations upon mutation.

#### 3.2.1 Convergence of the truncated channel ensemble

The channel partition function Z _*AB*_ sums over all simple paths connecting two residues in the spanning-tree ensemble. The number of self-avoiding paths grows exponentially with the graph size; the sum is therefore truncated at a maximum node length *L* = 9 (See Supplementary Material Section S1). The validity of this truncation is not a numerical convenience but a physical requirement: the channel thermodynamic functions are meaningful only if the truncated ensemble captures the essential conductance of the full network.

The convergence criterion for channel thermodynamic functions for a channel between residues a and b is provided by the dynamic distance *R*_*ab*_ given by Eq. 7. This quantity encodes the total informatics of the protein between the two coordinates, summed over all spanning trees and all path lengths, in a single matrix operation. It is the ground truth against which the truncation is measured. The subset dynamic resistance *R*_*ab*_ (*L*) is constructed as follows. For each maximum path length *L*, all simple paths of node length ≤ *L* are enumerated. The union of nodes visited by these paths defines a subset of the protein graph. The induced subgraph *G* (*L*) is formed by retaining every edge of *G* between any two visited nodes, not merely the edges that appear on the enumerated paths. This distinction is physically essential: a shortcut edge connecting two path-visited nodes provides a parallel information route whose contribution to the dynamic distance is not represented by any individual path in the truncated ensemble, yet whose removal would measurably increase the subset resistance to information flow. The dynamic distance *R*_*ab*_ (*L*) is computed from the Kirchhoff pseudoinverse of *G* (*L*), and the convergence ratio is *r* (*L*) = *R*_*ab*_ (*L*) / *R*_*ab*_ (*full*) . Rayleigh’s monotonicity principle [26] guarantees that the removal of edges from a resistive network cannot decrease the resistance. It follows that *R*_*ab*_ (*L*) ≥ *R*_*ab*_ (*full*), that the ratio *r* (*L*) 1, and that *r* (*L*) decreases monotonically toward unity as the induced subgraph approaches the full graph.

Figure 2 presents *R*_*ab*_ (*L*) for ten allosteric channels in the wild-type (6GOD) and G12D mutant (6GOF) ensembles. The exact resistance *R*_*ab*_ (*full*) is indicated by horizontal dashed lines. The decrease of *R*_*ab*_ (*L*) toward *R*_*ab*_ (*full*) reflects the progressive reconstruction of channel informatics: at each successive path length, new nodes are recruited into the induced subgraph, new edges are inherited from the full graph, and new parallel routes become available to the allosteric current.

**Figure 2.**
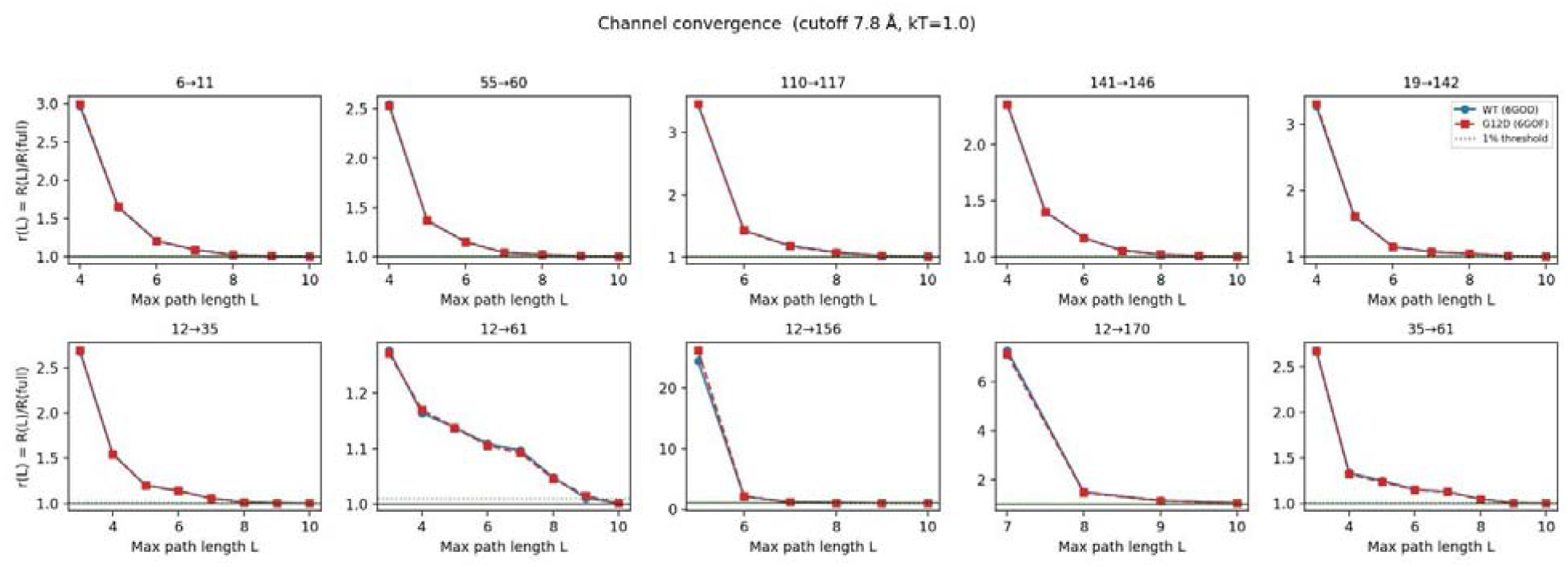
Convergence of the truncated channel dynamic distance for the ten allosteric channels in wild-type KRAS (6GOD, solid) and the G12D mutant (6GOF, dashed). Each panel shows the convergence ratio as a function of the maximum path length *L* (in nodes). The red dotted line marks the 99% threshold of the full protein. As an example, the P-loop channel (Res 6→11) reaches r ≥ 0.99 by *L* = 8, reflecting its compact local topology; the pre-Switch II (Res 55→60) and SAK (Res 141→146) channels reach r ≥ 0.98 by *L* = 9; the guanine base binding site (Res 110→117) and inter-lobe anchor (Res 19→142) channels reach r ≈ 0.93–0.97 at *L* = 9, indicating that their communication pathways sample a larger fraction of the protein edge set. Wild-type and G12D profiles are indistinguishable at this scale, confirming that the mutation scales the resistance uniformly across all path lengths rather than selectively reorganizing short- or long-range routes. All calculations were performed at kT = 1.0 Å with a contact cutoff of 7.8 Å.

Because the convergence analysis confirms that the truncated ensemble recovers between 93% and 99% of the exact dynamic distance in all channels, the thermodynamic functions so obtained are well-converged characterizations of the local allosteric landscape. The channel heat capacity C is the central quantity: it measures the breadth of the energy distribution sampled by the path ensemble connecting two residues, and its response to the G12D substitution shows whether the mutation stiffens, softens, or leaves unaltered each communication route. A large C indicates that the channel samples many energetically distinct path configurations; a small C signals that communication is confined to a narrow, energetically rigid subset of routes. The results are collected in Table 2 in the following section.

**Table 2.** Channel-resolved thermodynamic decomposition of allosteric heat capacity in wild-type and G12D KRAS. Channel heat capacities (*C*_WT_ and *C*_G12D_) evaluated at reference temperature *kT* = 1.0 over all enumerated paths of node length *L* ∈[2, 9] (contact cutoff *r*_*c*_ = 7.8*Å*). Total channel heat capacity change (Δ*C* = *C*_G12D_ − *C*_WT_) is decomposed into additive physical (Δ*C*_*E*_), topological (Δ*C*_*T*_), and cross-covariance (Δ*C*_*X*_) components (Δ*C* = Δ*C*_*E*_ + Δ*C*_*T*_ + Δ*C*_*X*_). Columns *n*_WT_ and *n*_G12D_ report the total number of accepted paths in the thermodynamic ensemble. PDB structures: WT = 6GOD; G12D = 6GOF.

| Channel | $C_{WT}$ | $C_{G12D}$ | $\Delta C$ | $\Delta C_E$ | $\Delta C_T$ | $\Delta C_X$ | $n_{WT}$ | $n_{G12D}$ |
| --- | --- | --- | --- | --- | --- | --- | --- | --- |
| Ch1 P-loop (6→11) | 6.7058 | 6.6512 | -0.0546 | -0.4610 | -0.1929 | +0.5993 | 946,282 | 968,240 |
| Ch2 pre-Switch II (55→60) | 6.2040 | 6.1345 | -0.0695 | -0.0510 | -0.0105 | -0.0080 | 638,606 | 642,205 |
| Ch3 GBS (110→117) | 8.0118 | 8.0468 | +0.0350 | +0.2612 | +0.1311 | -0.3574 | 824,774 | 858,454 |
| Ch4 SAK (141→146) | 6.0955 | 6.0897 | -0.0058 | -0.1788 | +0.0530 | +0.1199 | 934,827 | 964,794 |
| Ch5 Lobe linker (19→142) | 5.4593 | 5.3864 | -0.0729 | +0.2564 | +0.4943 | -0.8235 | 864,439 | 903,899 |
| Ch6 G12→Switch I (12→35) | 3.9708 | 4.0150 | +0.0442 | -0.2296 | -0.0961 | +0.3700 | 321,499 | 331,846 |
| Ch7 G12→Q61 SwII (12→61) | 2.0557 | 2.0485 | -0.0072 | +2.3445 | +2.4503 | -4.8021 | 124,222 | 134,205 |
| Ch8 G12→alpha5 (12→156) | 4.5710 | 4.6158 | +0.0448 | -0.6329 | -0.5001 | +1.1779 | 350,220 | 365,318 |
| Ch9 G12→C-term (12→170) | 3.7985 | 3.9351 | +0.1366 | +0.0267 | -0.0760 | +0.1859 | 12,372 | 14,003 |
| Ch10 Switch I→II (35→61) | 3.2064 | 3.2486 | +0.0421 | -0.5478 | -0.3986 | +0.9885 | 187,871 | 197,003 |

#### 3.2.2 Changes in channel thermodynamics upon mutation

To understand how mutational perturbation impacts the energetic and dynamic architecture of individual signaling corridors, we performed a channel-resolved decomposition of the heat capacity (*C*_*V*_) across all ten primary paths in wild-type and G12D KRAS **(Table 2)**. While global heat capacity remains robustly buffered (+0.50%), analyzing the localized channel sub-ensembles shows whether this stability stems from static structural rigidity or active thermodynamic compensation. By decomposing the total net shift (Δ*C*) into its constituent energetic (Δ*C*_*E*_), topological (Δ*C*_*T*_), and cross-covariance (Δ*C*_*X*_) terms, we can trace how internal entropic flux is redistributed across the network upon oncogenic mutation.

While net channel heat capacities (Δ*C*) remain remarkably stable, a channel-resolved decomposition across wild-type and G12D KRAS shows that this stability is maintained through strong internal thermodynamic compensation rather than structural rigidity (Table 2). Instead of a global collapse, the total heat capacity remains robustly buffered (+0.50%), while the underlying constituent terms, energetic (Δ*C*_*E*_), topological (Δ*C*_*T*_), and cross-channel covariance (Δ*C*_*X*_), undergo extensive shifts. Most notably, Channel 7 (G12 ^→^ Q61 / Switch II) features large positive energetic (Δ*C*_*E*_ = +2.345) and topological (Δ*C*_*T*_ = +2.450) shifts that are counterbalanced by negative cross-channel covariance (Δ*C*_*X*_ = −4.802). Furthermore, across every channel, the G12D mutation produces a consistent expansion in the effective microstate ensemble (*n*_G12D_ > *n*_WT_). This demonstrates that the mutation does not globally rigidify the protein, but rather redistributes the informational flux through delicate internal thermodynamic trade-offs across distinct functional sub-ensembles.

#### Thermodynamic Decomposition and the Cross-Coupling Compensation Principle

A prominent feature emerging from the decomposition of channel heat capacities (Table 2) is the synchronized co-variation of the energetic (*C*_*E*_) and topological (*C*_*T*_) components, balanced by an opposing shift in the cross-coupling term (*C*_*X*_). Across all evaluated functional routes, whenever a mutational perturbation causes *C*_*E*_ and *C*_*T*_ to shift in one direction, *C*_*X*_ systematically moves in the opposite direction.

Biophysically, *C*_*E*_ and *C*_*T*_ capture local contact rigidity and path multiplicity, respectively; a local geometric rearrangement that alters contact weights (*C*_*E*_) simultaneously constrains the local spanning-tree sub-ensemble (*C*_*T*_). The cross-coupling term *C*_*X*_, which quantifies the covariance between local path energies and global Kirchhoff matrix connectivity, acts as a thermodynamic buffer. By opposing the combined (*C*_*E*_ + *C*_*T*_) shift, *C*_*X*_ prevents local structural perturbations from destabilizing the total channel heat capacity (*C* = *C*_*E*_ + *C*_*T*_ + 2*C*_*X*_). This compensatory mechanism demonstrates that the surrounding protein matrix acts as an entropic reservoir, allowing KRAS to alter internal information routing (pathway rewiring) while maintaining robust, homeostatic channel capacity.

#### 3.2.3 Redistribution of path probabilities upon mutation

Jensen-Shannon Divergence *D*_*JS*_ indicates a negligible global shift in the statistical distribution of the path ensembles across most channels. This suggests that the G12D mutation does not degrade the informational capacity or the noise-profile of the communication, maintaining the channels as invariant conduits. However, this global stability masks a profound internal reorganization. While the volume of information remains constant, the specific routes through which it travels are fundamentally altered.

To map this internal shift, we calculate the allosteric importance *I*_*k*_ for each residue within the channel. This metric represents the fraction of the path ensemble that passes through a specific residue *k* . By comparing *I*_*k*_ between the wild-type and mutant, we identify residues that are recruited into the communication channel and those that are abandoned. Table 3 lists the residues that show the largest changes in importance, providing a direct map of the path re-partitioning caused by the G12D mutation.

**Table 3.** Channel Path Redistribution and Specific Allosteric Importance Shifts. Changes are given as percentages Δ*I*_*k*_. All ten channels are listed together, in a single ordered set.

| Channel | Residue-level allosteric importance shifts, $\Delta I_k$ (%) | | | | |
| --- | --- | --- | --- | --- | --- |
| **Res 6 → 11** (P-loop) | **G60** | **G77** | **G12** | **K16** | **V14** |
|  | +15.4 | +18.5 | -17.8 | +17.2 | +19.1 |
| **Res 55 → 60** (pre-SwII) | **S39** | **Q61** | **G12** | **L79** | **E62** |
|  | +15.4 | +22.7 | -25.9 | +18.8 | -17.8 |
| **Res 110 → 117** (GBS) | **S145** |  |  |  |  |
|  | +12.7 |  |  |  |  |
| **Res 141 → 146** (SAK) | **D154** | **V81** | **V112** | **G15** |  |
|  | -15.8 | +9.4 | +10.0 | +5.3 |  |
| **Res 19 → 142** (Lobe link) | **A155** | **D153** | **F156** | **C118** |  |
|  | +29.1 | +17.4 | +35.5 | -14.8 |  |
| **Res 12 → 35** (P-loop→SwI) | **E62** | **Q61** | **Y32** | **D33** |  |
|  | +19.3 | +18.6 | -15.9 | -14.9 |  |
| **Res 12 → 61** (canonical axis) | **G13** | **E62** |  |  |  |
|  | +15.3 | -12.6 |  |  |  |
| **Res 12 → 156** (P-loop→$\alpha 5$) | **I142** | **M111** | | | |
|  | +17.5 | -10.9 |  |  |  |
| **Res 12 → 170** (P-loop→C-term) | **G60** | **T58** | **K169** |  |  |
|  | +13.1 | +10.7 | -10.4 |  |  |

| Channel | Residue-level allosteric importance shifts, $\Delta I_k$ (%) | |
| --- | --- | --- |
|  | G13 | G12 |
| Res 35 → 61 (SwI→SwII) | -13.5 | -10.2 |

This table lists the residues *k* with the largest changes in allosteric importance *I*_*k*_ = *Z*_AB_(*k*)/*Z*_AB_, the fraction of the channel path ensemble passing through residue *k*, upon the G12D mutation. Changes are the relative shift 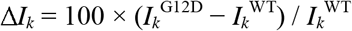 (positive = recruited, negative = abandoned; *I*_*k*_ is channel-specific, so a residue may shift in opposite directions in different channels); only residues with a wild-type occupancy *I*_k_ ^WT^ ≥ 3% are retained. All quantities were computed on *C*_α_ contact graphs (cutoff 7.8 Å, *kT* = 1.0 Å) for the wild-type (6GOD) and G12D (6GOF) structures, summing over all simple paths of node length ≤ 9. Despite the near-invariant channel heat capacity (Δ*C* ≈ 0, Table 2), these shifts show a precise re-partitioning of the information-transfer paths: the mutation abandons G12 and recruits Q61 and the inter-lobe bridge F156/A155, a pattern extended along the P-loop→Switch axes (recruitment of Q61 and I142; abandonment of G12 and of the Switch I residue Y32). The P-loop→C-terminal corridor 12→170 converges more slowly than the others, recovering ≈87% of the exact dynamic resistance at *L* = 9 (95% at *L* = 10); its shifts, carried by the flexible C-terminal residues G60/T58/K169, should be read as provisional at this truncation.

#### Detailed Biological and Biophysical Interpretation

1. **The Dominant Rewiring Hub at F156 (+35.5%):** The single largest importance surge across the entire network occurs at **F156 (+35.5%)** along the **Res 19** ^→^ **142 (Lobe link)** axis, anchored by **A155 (+29.1%)** and **D153 (+17.4%)**. Mechanistically, this suggests that the mutation drives a massive reorganization of allosteric information through the C-terminal *β* -sheet/ *α* -helix interface (Lobe 2 link). Rather than remaining confined to the immediate active site, signal transmission is heavily diverted toward the C-terminal structural anchor (*β* 7 /*α* 5 region), establishing F156 as a primary allosteric coordinate hub.
2. **Decoupling of Direct Switches and Amplification of the Catalytic Loop:** Along the Res 55 → 60 (pre-Switch II) channel we observe a large gain at Q61 (+22.7%) and at L79 (+18.8%), contrasted by strong suppression at G12 (−25.9%) and E62 (−17.8%). The canonical Res 12 → 61 axis is the most conservative channel in the set: only G13 (+15.3%) and E62 (−12.6%) exceed the reporting threshold, consistent with its near-zero ΔC (Table 2). The Res 35 → 61 (SwI→SwII) channel shows negative shifts at both P-loop glycines, G13 (−13.5%) and G12 (−10.2%), while the Res 12 → 35 (P-loop→SwI) channel abandons the Switch I residues Y32 (−15.9%) and D33 (−14.9%) in favour of Q61 (+18.6%) and E62 (+19.3%). Biological meaning: this is an allosteric decoupling effect. The mutation weakens direct P-loop↔Switch I and Switch I↔Switch II cross-talk negative Δ*I*_*k*_ at G12, G13, Y32 and D33 across the 12→35 and 35→61 channels and reroutes communication through an amplified Q61/E62 Switch II axis.
3. **Long-Range Inter-Lobe Relay (P-loop to C-terminus):** The Res 12 → 156 channel shows a substantial gain at I142 (+17.5%) with a compensating loss at M111 (−10.9%). The Res 12 → 170 corridor recruits G60 (+13.1%) and T58 (+10.7%) at the expense of K169 (−10.4%). Biological meaning: the active site establishes long-range energetic pathways extending to the C-terminal region. Taken together with the recruitment of Q61 and E62 one channel upstream (Res 12 → 35), the gain at I142 indicates that Switch II and the β6/β7 region act as relay stations bridging active-site dynamics to the lower lobe. Because 12→170 is the least converged of the ten channels (Figure 2), the shifts listed for it are provisional.
4. **Guanine Binding Site (GBS) Localized Adjustment:** In the Res 110 → 117 (GBS) channel the only shift above the reporting threshold is a gain at S145 (+12.7%), so the guanine-recognition corridor is essentially invariant under the mutation. The adjacent SAK channel (Res 141 → 146) is more responsive: it abandons D154 (−15.8%) and redistributes traffic to the hydrophobic core residues V112 (+10.0%) and V81 (+9.4%) and to G15 (+5.3%). Biological meaning: the nucleotide-binding channels undergo only localized entropic redistribution, shifting communication duties onto adjacent hydrophobic core residues while preserving structural integrity around the guanine ring.

## 4. Discussion

### 4.1 A spanning-tree partition function as the natural language of allostery

The central result of this work is both conceptual and computational: allosteric communication can be placed on the same footing as classical statistical mechanics without invoking molecular dynamics trajectories, normal-mode expansions, or empirically trained potentials. The spanning tree, the minimal cycle-free subgraph that preserves global connectivity, is the natural microstate of a communication network precisely because it contains exactly one unambiguous route between any two nodes. Identifying these microstates with Boltzmann weights derived from inter-residue distances, and summing their contributions via Kirchhoff’s Matrix-Tree Theorem, collapses an exponentially large state space into a single determinantal evaluation. The resulting partition function is not an approximation; it is exact for any contact graph and any positive edge weights.

This exactness has a practical corollary: every thermodynamic quantity derived here; free energy, entropy, heat capacity, and all channel analogs, is guaranteed to be consistent and self-contained within the chosen graph representation. There are no uncontrolled approximations; any discrepancy between this framework and experiment reflects the suitability of the graph model (choice of cutoff, weight function, or structural ensemble), not the statistical mechanical machinery. This separability of modeling choices from theoretical rigor is a key advantage over perturbative or simulation-based approaches, where numerical and sampling errors are inseparable from the physics.

### 4.2 Preservation of global thermodynamics without structural reorganization

While conventional structural metrics, such as root-mean-square deviation (RMSD) comparisons, contact-map overlays, and standard molecular dynamics observables, show that wild-type and mutant KRAS share a nearly identical mean structure, our topological framework confirms that global thermodynamic averages (internal energy, entropy, and global heat capacity) remain essentially unchanged upon the G12D substitution (< 0.5% variation).

In conventional thermodynamics, heat capacity measures a system’s ability to absorb energy through fluctuations, corresponding in our model to the variance in spanning-tree energies across accessible acyclic communication configurations. Rather than driving a macroscopic thermodynamic shift or global structural reorganization, the G12D mutation preserves these global averages while altering the internal routing of allosteric information.

This preservation of global thermodynamic metrics aligns with the Cooper-Dryden framework of allostery without conformational change [2]. However, our model extends this concept: rather than relying on harmonic vibrational entropy, the spanning-tree ensemble captures the combinatorial connectivity of the contact network. The primary impact of the G12D mutation is therefore not a global thermodynamic collapse, but a highly localized, channel-specific redistribution of information traffic across intermediate residues.

### 4.3 Invariant channel capacity with internal redistribution: a topological compensation principle

The remarkable finding of this analysis is the near-perfect conservation of channel-level heat capacities across the mutation, mirroring the preservation of global thermodynamic metrics. For each of the channels examined, | Δ*C* | is at most 0.08 in reduced units (less than 1.5% of the channel value). This shows a structural principle that may be called topological compensation: the G12D substitution preserves the overall variance of path energies within functionally critical channels while reorganizing internal routing.

This compensation is not passive. Decomposing each channel heat capacity into its energetic (*C*_*E*_), topological (*C*_*T*_), and cross-coupling (*C*_*X*_) components shows large, opposing shifts that cancel at the total level. For example, in the inter-lobe bridge channel (residues 19–142), the energetic component increases by +0.26, the topological component increases by +0.49, and the cross-term decreases by −0.82, yielding a net change of only −0.07 . These compensatory shifts reflect the mathematical coupling between the local path ensemble and the global Laplacian. The surrounding protein acts as a topological reservoir whose local modifications by the mutation precisely rebalance the channel’s internal thermodynamic components while preserving the total.

This topological compensation has a crucial functional implication: the G12D mutation does not destroy allosteric communication, but reorganizes it in a thermodynamically self-consistent manner. The channel retains its total information bandwidth while redirecting traffic through different residue intermediaries.

### 4.4 Residue-level redistribution and the oncogenic mechanism of G12D

The near-invariant Jensen–Shannon divergences confirm that the G12D mutation does not change the statistical character of the allosteric channels, it does not make them noisier or less informative. What it alters, precisely and quantitatively, is the identity of the residues carrying the allosteric current (Table 3). The abandonment of G12 as an allosteric hub (−17.8% in the P-loop channel, −25.9% in the pre-Switch II channel) is a direct geometric consequence of the glycine-to-aspartate substitution. Glycine’s minimal side chain makes it a flexible node in the contact network. Substituting it with aspartate alters local geometry, modifies the weight of adjacent edges in the Laplacian, and removes G12 from the dominant paths. The network does not break; it detours.

The recruitment of Q61 (+22.7% in the pre-Switch II channel) is particularly significant. Glutamine 61 is the catalytic residue responsible for activating the water molecule that executes GTP hydrolysis. That the G12D mutation increases Q61’s allosteric importance without mutating Q61 itself suggests a subtle mechanism: G12D redirects the allosteric current so that Q61 becomes a primary conduit for information transfer rather than a catalytic effector. This topological repurposing of Q61 may contribute to the sustained GTP-bound state of oncogenic KRAS by diverting Q61 from catalysis to communication. The dramatic shifts in the inter-lobe bridge channel (A155 +29.1%, F156 +35.5%, C118 −14.8%) are equally consequential. Residues A155 and F156 lie within the α5 helix of the allosteric lobe, at the C-terminal to the β6 strand; their increased importance concentrates inter-lobe communication through a narrow hydrophobic bridge rather than distributing it across the broader network. This localized inter-lobe traffic makes the two-domain architecture less accommodating to the conformational dynamics required for nucleotide exchange, providing a mechanistic basis for the altered exchange kinetics of G12D KRAS.

### 4.5 The principle of minimum surprisal and mutational hijacking of allosteric highways

The principle of minimum surprisal provides a unifying framework for these observations. In the wild-type protein, the signaling ensemble routes information through paths that balance energetic efficiency (short contacts) and topological redundancy. The G12D mutation reshapes this surprisal landscape: paths passing through the altered G12 node become less favorable, while alternative paths through Q61, A155, and F156 become comparatively more accessible.

This surprisal redistribution represents the mutational hijacking of the protein’s natural allosteric highways. The total information capacity of the channels is preserved, but specific routes prefer paths through residues whose functional roles make this redirection pathologically favorable. Q61 becomes an allosteric node rather than a catalytic one, A155 and F156 form a local bottleneck, and G12 is bypassed. The oncogenic state is thus not one of communication failure, but of communication capture.

### 4.6 Therapeutic implications: targeting topological bottlenecks

These findings offer concrete implications for designing allosteric KRAS inhibitors. Standard strategies targeting the GDP pocket or Switch I/II interface rely on mean-structure properties. Our analysis suggests that residues exhibiting large allosteric importance shifts upon mutation, specifically Q61 in the pre-Switch II channel and A155/F156 in the inter-lobe bridge, represent alternative targets. A ligand designed to selectively reduce the allosteric occupancy of these mutant-recruited hubs could restore wild-type traffic patterns without needing to displace the nucleotide or directly contact the G12 mutation site.

### 4.7 Scope, limitations, and future directions

Several methodological considerations are worth noting:

1. Static Structural Inputs: The analysis operates on single crystal structures (*C*_*α*_ backbones) rather than dynamic conformational ensembles. Incorporating molecular dynamics trajectories or normal-mode sampling would allow edge weights to be averaged over thermal distributions.
2. Parameter Sensitivity: The contact cutoff (*r*_*c*_ = 7.8*Å*), thermal parameter (*kT* = 1.0), and path truncation (*L*_max_ = 6) represent defined model inputs. Sensitivity analysis across parameter space (Supplementary Note S5, Figure S5.1) confirms that channel trends remain stable across parameter variations.
3. Path Truncation: Truncating path expansion at *L*_max_ = 6 captures the vast majority of information flux as an empirical observation. While capturing the dominant path probabilities, higher-order thermodynamic functions weight longer path lengths differently, making extended path calculations a natural direction for future refined models.
4. Coarse-Grained Geometry: Using *C*_*α*_ backbone coordinates senses mutations through local geometric shifts rather than explicit side-chain chemistry. Thus, residue-level importance shifts serve as predictive hypotheses for experimental or all-atom MD validation.

### 4.8 Relationship to existing frameworks

The Spanning-Tree Thermodynamics (STT) framework occupies a distinct position relative to traditional network models. In the high-temperature limit (*kT* →∞), all edge weights equalize and the partition function reduces to the number of spanning trees in an unweighted graph, recovering the Gaussian Network Model (GNM) as a special topological case. At finite temperature, Boltzmann weighting introduces a geometric bias favoring short, strong contacts, providing a physically motivated generalization without losing analytical tractability.

Unlike Monod-Wyman-Changeux models, STT requires no pre-specified discrete conformational states; microstates emerge naturally from graph topology. Unlike classical Cooper-Dryden allostery, which focuses on vibrational entropy, STT evaluates the combinatorial entropy of the spanning-tree ensemble. The Burton-Pemantle theorem serves as the exact bridge between the global network and local channels via the Moore-Penrose pseudoinverse of the Laplacian.

Finally, a fundamental statistical mechanical distinction separates STT from classical elastic models like GNM. While GNM operates under a strictly harmonic assumption yielding a constant, temperature-independent heat capacity, STT evaluates a discrete, combinatorial ensemble with a bounded spectrum. As temperature varies, this discrete ensemble exhibits a clear heat capacity maximum, a topological Schottky anomaly, characterizing the transition from a constrained ground topology to a degenerate network manifold (Supplementary Figure S3.1). This confirms that STT captures non-harmonic topological flexibility and entropic feedback mechanisms beyond the reach of standard harmonic models.

## Notes

https://github.com/fatmasenguler/Mutation_KRAS_analysis/tree/main

### Competing Interest Statement

The authors have declared no competing interest.

### Summary of Updates

This version of the manuscript has been thoroughly revised in response to the constructive feedback from both reviewers. During the revision process, we identified that the previously reported global decrease in heat capacity was an artifact caused by numerical instability in our initial calculations. To address this, we implemented a more robust computational methodology based on Laplacian trace identities. This technical improvement refined our results and led to a strategic refocusing of our manuscript. Rather than concentrating on broad changes in global thermodynamic properties, our updated analysis centers in detail on the localized rewiring of individual allosteric channels driven by the G12D mutation in KRAS. We have updated all calculations, revised the corresponding figures and tables, and expanded our discussion of channel-resolved allosteric communication. These modifications significantly improve the numerical accuracy, physical clarity, and overall scientific rigor of the manuscript.

