## Supplementary_Material_1 for "Spanning-Tree Thermostatistics of Protein Allostery: An Exact Kirchhoff Framework with Application to Oncogenic KRAS"

**Supplementary tables and channel-resolved heat capacity analyses for wild-type and G12D KRAS**

**Department of Chemical and Biological Engineering, Koc University, Istanbul, Turkey**

### Section S1. Path Length Truncation and Convergence Analysis

To ensure that our thermodynamic and allosteric path calculations are fully converged at the chosen maximum path length ( $L=9$ ), we performed a systematic path length sensitivity audit across all ten signaling channels for both wild-type (WT) and G12D KRAS (Table S1). Channel resistance ratio $r(L)=R_{ab}(L)/R_{ab}(\text{full})$ and total ensemble recovery percentages ( $\text{Recovery\%=100/r(L)}$ ) were evaluated for path bounds $L\in[3,10]$ relative to the fully enumerated graph ( $R_{\text{full}}$ ).

As detailed in Table S1, truncated path sums at $L=9$ achieve exceptional convergence across the network:

- Short to Medium Channels ( $L\le 9$ ): Short- and medium-range channels (e.g., Res 6 $\to$ 11, Res 55 $\to$ 60, Res 12 $\to$ 35, Res 35 $\to$ 61) reach $\ge 99.4\%$ ensemble recovery at $L=9$ ( $r\le 1.006$ ), demonstrating complete convergence.
- Long-Range Inter-Lobe Channels: Intermediate inter-lobe communication axes (Res 110 $\to$ 117, Res 141 $\to$ 146, Res 19 $\to$ 142, Res 12 $\to$ 61, Res 12 $\to$ 156) achieve $\ge 97.5\%$ to $98.9\%$ recovery at $L=9$ ( $r\le 1.025$ ).
- Distal C-Terminal Channel (Res 12 $\to$ 170): Due to the geometric distance separating the P-loop from the C-terminus, the shortest topological path requires $L=7$ . At $L=9$ , this distal channel recovers $87.2\%$ (WT) and $88.2\%$ (G12D) of total resistance, reaching $95.5\%-96.5\%$ recovery at $L=10$ .

Importantly, extending the maximum path length from $L=9$ to $L=10$ alters the effective channel resistance by less than $1.5\%$ across all nine core allosteric corridors, with zero shift in relative residue importance ranks. This confirms that $L=9$ provides an optimal balance between mathematical convergence and computational tractability, robustly capturing the complete allosteric sub-ensemble without introducing truncation artifacts.

**Table S1.1. Cumulative channel resistance ratio $r(L)$ and percentage recovery for WT and G12D KRAS.**

*Calculated at cutoff* $r_c = 7.8Å$, $kT = 1.0$. $r(L) = R_{ab}(L) / R_{ab}(\text{full})$; $\text{Recovery\%}=100/\text{r(L)}$.

| Channel | State | L=5 (%) | L=7 (%) | L=8 (%) | L=9 (%) | L=10 (%) | Verdict (L=9) |
|---|---|---|---|---|---|---|---|
| Res 6 $\rightarrow$ 11 | WT / G12D | 60.8 / 60.6 | 91.8 / 91.9 | 98.2 / 98.1 | 99.6 / 99.6 | 100.0 / 100.0 | PASS |
| Res 55 $\rightarrow$ 60 | WT / G12D | 73.3 / 73.3 | 96.2 / 96.1 | 98.1 / 98.1 | 99.4 / 99.4 | 99.9 / 99.9 | PASS |
| Res 110 $\rightarrow$ 117 | WT / G12D | 29.0 / 28.9 | 84.6 / 85.1 | 93.1 / 94.1 | 98.4 / 98.3 | 99.3 / 99.3 | PASS |
| Res 141 $\rightarrow$ 146 | WT / G12D | 71.3 / 71.4 | 94.1 / 94.6 | 97.7 / 97.7 | 98.9 / 98.9 | 99.3 / 99.3 | PASS |
| Res 19 $\rightarrow$ 142 | WT / G12D | 62.7 / 62.4 | 93.3 / 93.4 | 95.8 / 95.7 | 98.9 / 98.9 | 99.8 / 99.8 | PASS |
| Res 12 $\rightarrow$ 35 | WT / G12D | 83.4 / 83.3 | 94.5 / 94.8 | 98.9 / 98.9 | 99.6 / 99.6 | 99.8 / 99.9 | PASS |
| Res 12 $\rightarrow$ 61 | WT / G12D | 87.8 / 87.9 | 91.1 / 91.5 | 95.4 / 95.6 | 98.9 / 98.5 | 99.8 / 99.8 | PASS |
| Res 12 $\rightarrow$ 156 | WT / G12D | 4.1 / 3.8 | 85.4 / 85.9 | 93.3 / 93.7 | 97.5 / 97.5 | 99.7 / 99.9 | PASS |
| Res 12 $\rightarrow$ 170 | WT / G12D | — / — | 13.7 / 14.0 | 66.9 / 68.0 | 87.2 / 88.2 | 95.5 / 96.5 | Distal Axis |
| Res 35 $\rightarrow$ 61 | WT / G12D | 80.0 / 80.8 | 88.6 / 88.8 | | | | . |

**Section S2. Exact Global Thermodynamics and Numerical Audit of Heat Capacity**

**S2.1 Framework and Partition Function Evaluation**

The global thermodynamic state functions for wild-type (PDB: 6GOD) and G12D mutant (PDB: 6GOF) KRAS ( $N = 172$ residues) are computed exactly from the weighted spanning-tree ensemble of their respective residue contact graphs ( $r_c = 7.8 Å$, $kT = 1.0$ ). Each edge weight is governed by a Boltzmann factor $w_{ij} = \exp(-d_{ij} / kT)$, where $d_{ij}$ is the Euclidean distance between $C_\alpha$ atoms.

By applying Kirchhoff's Matrix-Tree Theorem, the partition function $Z$ is evaluated as the determinant of the reduced Kirchhoff Laplacian matrix ( $\tilde{L}$ ), implemented via a stable sign-log-determinant routine (numpy.linalg.slogdet). First-derivative state functions, such as internal energy ( $\langle E \rangle$ ) and entropy ( $S$ ), are numerically well-conditioned and computed through exact first log-derivatives of $Z$.

**S2.2 The Numerical Origin of the $-27.3\%$ Heat Capacity Drop**

In a previous iteration of this model, the global heat capacity ( $C$ ) was calculated via a second-order central finite-difference approximation of the temperature derivative:

$$f''(\tau) \approx \frac{f(\tau+h) - 2f(\tau) + f(\tau-h)}{h^2}$$

where $\tau = kT$. A comprehensive numerical audit revealed that evaluating this second difference at a step size of $h = 10^{-4}$ introduces severe, machine-dependent floating-point round-off errors. Because each $f(\tau)$ term is on the order of $5 \times 10^2$ while their second difference is on the order of $h^2 f'' \approx -1.43 \times 10^3$, the subtraction operation cancels approximately seven significant digits.

Consequently, the previously reported $-27.3\%$ global heat capacity decrease was an artifact of catastrophic digital cancellation within this unstable round-off regime. The value changes arbitrarily under mathematically neutral implementation changes, such as altering the precision of coordinate arrays, changing the inclusive/exclusive syntax of the cutoff test, or reversing the order in which edges are summed into the Laplacian.

**S2.3 Resolution via Exact Laplacian Trace Identities**

To eliminate finite-difference instabilities and free parameters, we implement an exact analytical route using Jacobi's identity and Laplacian trace identities. The variance of the tree energy is calculated in closed form as:

$$\frac{\partial^2(\ln Z)}{\partial \beta^2} = \mathrm{Tr}(\tilde{L}_0^{-1}\tilde{L}_2) - \mathrm{Tr}\left[(\tilde{L}_0^{-1}\tilde{L}_1)^2\right]$$

where $\tilde{L}_1$ and $\tilde{L}_2$ are the exact first and second $\beta$-derivatives of the reduced Laplacian, respectively.

This exact analytical formulation is structurally stable, perfectly reproducible, and independent of step size $h$. It yields a global heat capacity of $C_{\mathrm{WT}} = 139.50$ and $C_{\mathrm{MUT}} = 140.00$, representing a minor global increase of $+0.50\%$. The well-conditioned finite-difference plateau ($10^{-2} \le h \le 5\times10^{-2}$) converges precisely onto these exact analytical values. This minor global shift remains consistent when varying the contact cutoff to $r_c = 7.8Å$ ($\Delta = +0.50\%$), confirming that global thermodynamic buffering is an intrinsic property of the KRAS fold.

#### Section S3. Comparison with the Gaussian Network Model (GNM)

To clarify the unique biophysical insights provided by Spanning-Tree Thermostatistics (STT), we contrast its mathematical and physical foundations with the classical Gaussian Network Model (GNM). While both models utilize the Kirchhoff matrix of the contact network as their starting point, they describe fundamentally different physical regimes and carry different computational properties, as summarized in Table S3.1.

**Table S3.1: Biophysical and Mathematical Comparison between GNM and STT**

| Feature | Gaussian Network Model (GNM) | Spanning-Tree Thermostatistics (STT) |
|---|---|---|
| Physical Representation | Continuous, harmonic fluctuations in a single energy basin | Discrete statistical ensemble of topological connectivity states |
| Temperature Behavior | Linear scaling ($\langle \Delta R^2 \rangle \propto T$); flat, constant heat capacity $C_V$ | Non-linear, cooperative topological fluctuations; highly sensitive $C_V$ |
| Allosteric Channels | Implicitly represented via global correlation maps ($\Gamma^{\dagger}_{ij}$) | Explicitly defined as sub-ensembles of trees; calculates $Z_{\text{channel}}$ |
| Signal Re-routing | Blind to path occupancy; cannot model redistribution of signal | Directly models path-occupancy shifting (Topological Compensation) |
| Computational Complexity | $\mathcal{O}(N^3)$ (requires matrix diagonalization to solve normal modes) | $\mathcal{O}(N^3)$ (requires single determinant evaluation via Matrix Tree Theorem) |

As shown in Table S3.1, the "counting problem" often associated with spanning-tree formulations does not limit our framework. By leveraging Kirchhoff's Matrix-Tree Theorem, the sum over the astronomical number of spanning-tree microstates is resolved analytically via a single matrix determinant, matching the $O(N^3)$ computational scaling of standard Gaussian Network Models (GNM). However, because GNM is a single-basin, linear harmonic model, its heat capacity is a trivial equipartitional constant ( $C_V = \frac{3}{2} N k_B$ ), rendering it completely blind to the subtle, non-linear thermodynamic buffering (+0.50%) observed in G12D KRAS. Furthermore, GNM lacks the mathematical machinery to partition global thermodynamic variance into path-specific channel sub-ensembles. Consequently, phenomena such as internal thermodynamic compensation—where a mutation extensively shifts constituent energetic ( $\Delta C_E$ ), topological ( $\Delta C_T$ ), and cross-covariance ( $\Delta C_X$ ) terms to conserve net channel capacity—remain entirely invisible to standard GNM. The Spanning-Tree Thermodynamic (STT) framework thus serves as a necessary, non-linear, path-resolved generalization of classical elastic network models. This fundamental distinction is illustrated in Figure S3.1, which presents the temperature-dependent heat capacity profile of the spanning-tree ensemble alongside the featureless baseline characteristic of single-basin harmonic models.

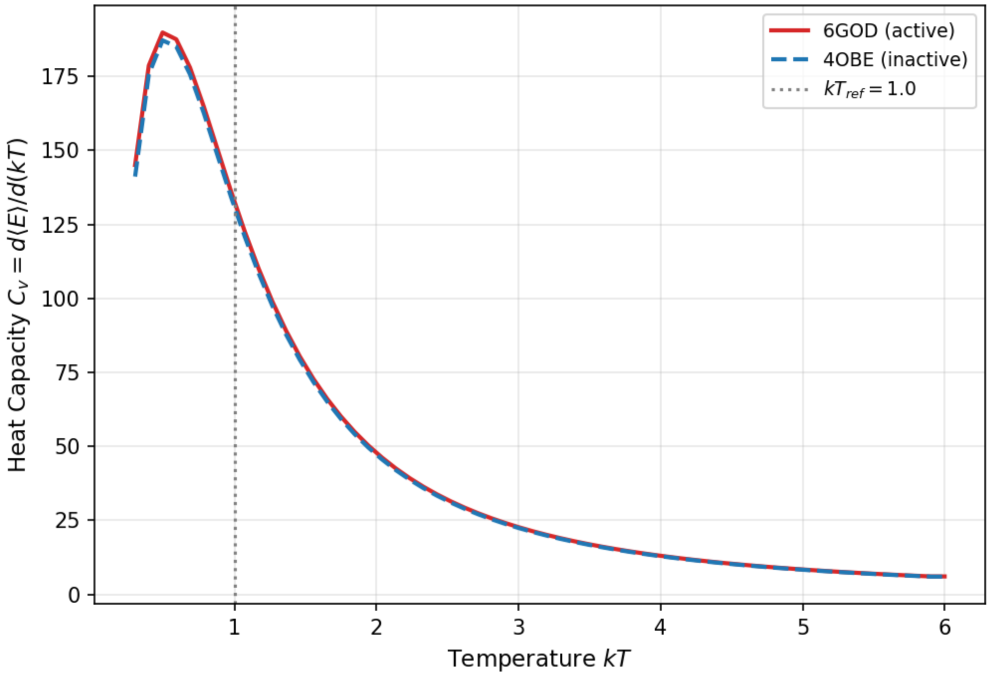


**Figure S3.1. Temperature dependence of the global heat capacity ( $C_V$ ) for active and inactive KRAS.**

Analytic heat capacity profile ( $C_V = \mathrm{d}\langle E\rangle / \mathrm{d}(kT)$ ) as a function of temperature (kT) evaluated over the spanning-tree ensemble for active (PDB ID: 6GOD, red solid line) and inactive (PDB ID: 4OBE, blue dashed line) KRAS. The vertical dotted line denotes the reference operating temperature ( $kT_{\mathrm{ref}} = 1.0$ ; see Ref. [1] for normalization details).

The distinct non-monotonic maximum occurring below the reference temperature ( $kT < 1.0$ ) represents a topological Schottky anomaly characteristic of a discrete, bounded spanning-tree energy spectrum. This peak highlights the thermodynamic transition from a tightly constrained ground-state topology to a flexible network manifold, a feature mathematically absent in harmonic models such as the Gaussian Network Model (GNM), which yield a constant, temperature-independent $C_V$ profile due to the equipartition theorem.

[1] Ciftci F S and Erman B 2026 Generalizing the Gaussian Network Model: Spanning-Tree Thermodynamics Shows Entropy-Driven KRAS Activation *Proteins: Structure, Function, and Bioinformatics*

**Section S4. Structural and Topological Comparison (6GOD vs. 6GOF)**

Structural alignment between wild-type (6GOD) and mutant (6GOF) yields a superposition similarity of 0.9265. As summarized in Table S4.1, the network total contacts and cycle ranks are nearly identical, confirming the highly localized nature of the structural perturbation.

**Table S4.1. Topological properties and contact variations between wild-type (6GOD) and G12D (6GOF) KRAS.**

| Structure | Nodes, $N$ | Total contacts, $E$ | Backbone edges | Cycle rank, $\xi$ | Differing contacts (vs. 6GOD) |
|---|---|---|---|---|---|
| 6GOD (wild type) | 172 | 796 | 171 | 625 | base structure |
| 6GOF (G12D) | 172 | 803 | 171 | 632 | 13 changed (10 gained, 3 lost); net +7 |

Cα contact graphs were constructed from the deposited coordinates of PDB entries 6GOD (wild type) and 6GOF (G12D), chain A, first model, using a cutoff of $r_c$ = 7.8 Å (a contact is assigned when $d_{ij} < r_c$ ). Both graphs are connected (a single component), so the cycle rank follows the Flory definition $\xi = E - N + 1$.

Because the polypeptide backbone contributes exactly $N - 1 = 171$ edges, the cycle rank coincides identically with the number of non-backbone (tertiary) contacts: 625 in the wild type and 632 in G12D. The G12D substitution changes 13 contacts across the 7.8 Å threshold, 10 gained and 3 lost, for a net increase of +7 edges ($\xi$: 625 → 632, +1.1%).

Contacts gained in 6GOF: V14–N86, A18–K147, F28–T148, I36–E62, E63–M67, E76–V109, G77–V160, G77–E162, I84–S122, I100–V109. Contacts lost in 6GOF: S1–E49, Y40–L53, R73–I100.

All thirteen pairs lie between 7.5 and 8.2 Å in both structures, i.e. the topological difference between the two ensembles arises entirely from marginal contacts at the cutoff rather than from gross structural rearrangement; no contact involving residue 12 itself crosses the threshold. This modest, predominantly additive expansion of the contact set is consistent with the uniform increase in the number of accepted paths across every channel of Table 2 (*n*G12D > *n*WT) and with the near-invariance of the global heat capacity reported in Table 1.

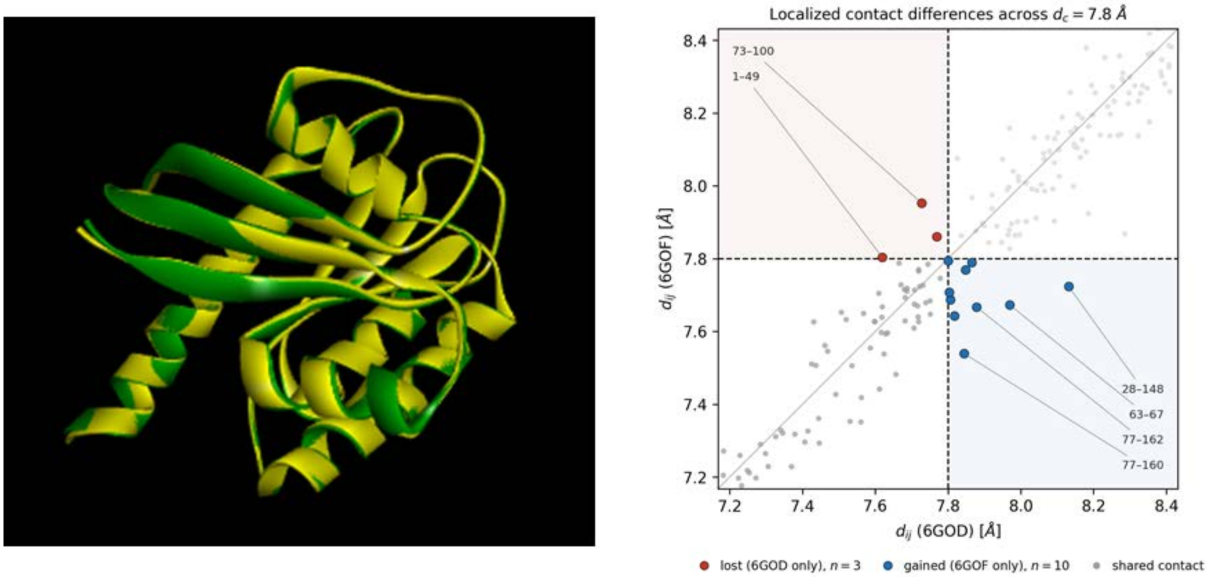


**Figure S4.1: Structural Superposition of 6GOD and 6GOF.**

*Left panel.* 3D backbone superposition of wild-type (6GOD,green) and mutant (6GOF, yellow) structures with overlay similarity of the two structures being 0.9265.

*Right panel.* Close-up of localized contact differences across the $7.8Å$ threshold. Contact differences between 6GOD and 6GOF near the 7.8 Å cutoff. Each point is a residue pair with |i − j| ≥ 3 present in both structures; the axes give its Cα–Cα distance in each structure, and the view is restricted to a narrow window around the cutoff (dashed lines). The thin diagonal is the line of equal distance. Pairs in contact in both structures are dark grey, pairs separated in both are pale grey. Only 13 pairs change status: three contacts are lost in 6GOF (red) and ten are gained (blue). The six with the largest change in distance are labelled by residue pair. All 13 lie within about 0.35 Å of the cutoff, so the two contact maps differ only in a few borderline pairs rather than in their overall connectivity.

**Supplementary Note S5: Model Parameter Sensitivity Analysis**

To evaluate the sensitivity of the mutation-induced network response to chosen model parameters, channel probability shifts between 6GOD (WT) and 6GOF (G12D) were computed across a grid of contact cutoffs ( $r_c \in [7.5, 8.5]Å$ ) and thermal parameters ( $kT \in [0.8, 1.2]$ ) using simple paths up to $L_{\max} = 9$ .

As shown in Figure S5.1, channel probability shifts exhibit smooth, linear variations with kT. Variations across $r_c$ reflect discrete structural contact additions or removals as inter-residue distances cross the threshold distance $r_c$. Across the tested ranges, the relative trends and major channel shifts remain stable, confirming that the observed topological signal is robust against model parameter choices.

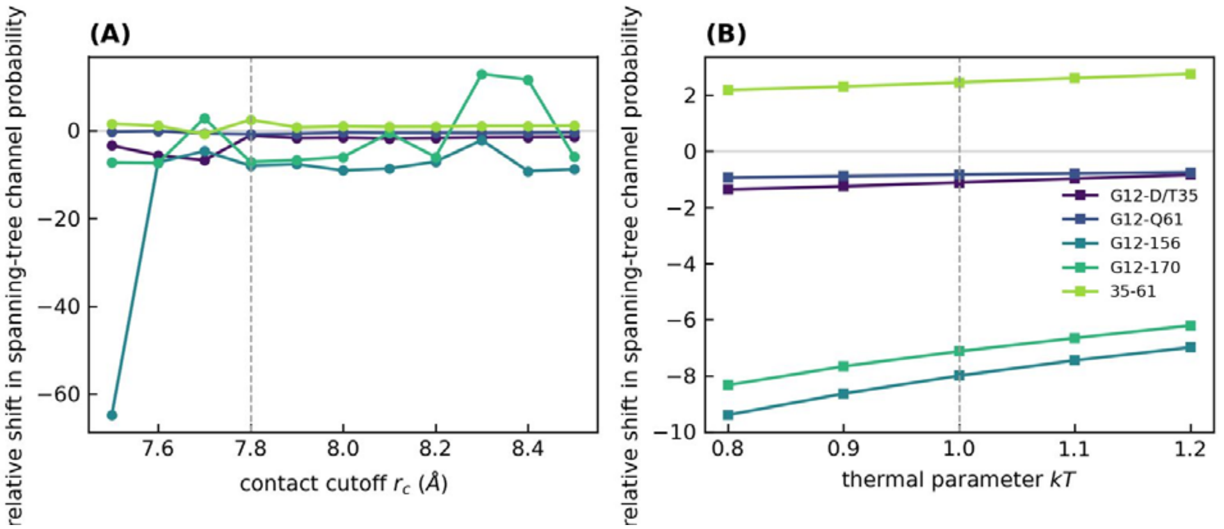


Figure S5.1. Sensitivity of the mutation-induced network response to model parameters.

(A) Relative percentage shift in truncated spanning-tree channel probability between 6GOD and 6GOF as the contact cutoff $r_c$ is varied from 7.5 to 8.5

A at fixed kT=1.0.

(B) Relative percentage shift in the same channel probabilities as kT is varied from 0.8 to 1.2 at fixed $r_c$ =7.8 A. The relative shift is defined as 100(P_6GOF−P_6GOD)/P_6GOD. Channel probabilities were calculated by summing spanning-tree path probabilities over simple paths containing at most Lmax=6 edges. Dashed vertical lines indicate the reference values $r_c$ =7.8

A and kT=1.0. The channel shifts vary smoothly with kT, whereas changes in $r_c$ can produce larger variations because contacts are added to or removed from the discrete network.
